# Isolation by Distance in Populations with Power-law Dispersal

**DOI:** 10.1101/2020.06.24.168211

**Authors:** Tyler B. Smith, Daniel B. Weissman

## Abstract

Limited dispersal of individuals between generations results in isolation by distance, in which individuals further apart in space tend to be less related. Classic models of isolation by distance assume that dispersal distances are drawn from a thin-tailed distribution and predict that the proportion of the genome that is identical by descent between a pair of individuals should decrease exponentially with the spatial separation between them. However, in many natural populations, individuals occasionally disperse over very long distances. In this work, we use mathematical analysis and coalescent simulations to study the effect of long-range (power-law) dispersal on patterns of isolation by distance. We find that it leads to power-law decay of identity-by-descent at large distances with the same exponent as dispersal. We also find that broad power-law dispersal produces another, shallow power-law decay of identity-by-descent at short distances. These results suggest that the distribution of long-range dispersal events could be estimated from sequencing large population samples taken from a wide range of spatial scales.

## Introduction

Populations with limited dispersal exhibit “isolation by distance”: the more distant individuals are from each other in space, the less related they tend to be (Wright 1946; Rohlf and Schnell 1971; Slatkin 1991). The strength of isolation by distance can be used to infer demography and dispersal (Koenig *et al*. 1996; Cayuela *et al*. 2018; Bradburd and Ralph 2019; Battey *et al*. 2020). For populations spread over a fairly continuous range, rather than being clumped into a small number of discrete sub-populations, dispersal is often assumed to be thin-tailed, with displacement approximately following a normal distribution (Barton *et al*. 2002; Ringbauer *et al*. 2017). If dispersal is unbiased and homogeneous, it is then characterized by a single parameter, the dispersal rate *D*. If we track the spatial position of a lineage backwards in time over multiple generations, its motion will approach a diffusion, with *D* as diffusion constant. While in general the spatial pattern even of neutral genetic diversity depends on selection (Barton *et al*. 2013a; Allman and Weissman 2018), for populations evolving completely neutrally the strength of isolation by distance is simply determined by the balance between dispersal and mutation. Specifically, pairwise genetic similarity is predicted to decay exponentially with distance, with a decay rate of 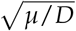, where *μ* is the mutation rate (Kimura and Weiss 1964; Malécot 1975; Slatkin and Barton 1989; Slatkin and Arter 1991; Slatkin 1993; Rousset 1997, 2000; Barton *et al*. 2002; Ralph and Coop 2013).

In many populations, however, dispersal is heavy-tailed, with individuals occasionally moving much farther than the typical distance (Willson 1993; Clark 1998; Atkinson *et al*. 2002; Baguette 2003; Brockmann *et al*. 2006; Dai *et al*. 2007; Fric and Konvicka 2007; Devaux *et al*. 2007; Aguillon *et al*. 2017; Vallaeys *et al*. 2017). In many such populations, the tail of the dispersal distribution can be approximated by a power law, i.e., the probability that an individual disperses farther than a long distance *y* in their lifetime is proportional to *y*^−*α*^ for some *α* > 0. The smaller *α*, the longer the tail; distributions with *α* ≤ 2 have infinite standard deviation, while those with *α* ≤ 1 also have infinite mean. (Of course, dispersal tails must be cut off at some distance corresponding to the diameter of the population’s range, and correspondingly when we say that the standard deviation or mean are “infinite” we really mean that they are set by the total range size.) These power-law dispersal distributions can lead to qualitatively different lineage trajectories from the ones produced by thin-tailed dispersal, especially if *α* < 2 (Metzler *et al*. 2009); see Fig. 1. In particular, for any *α*, the most likely way for a lineage to travel an unusually long distance over many generations is for it cover most of the distance in a single very long jump in one generation (Vezzani *et al*. 2019).

**Figure 1.**
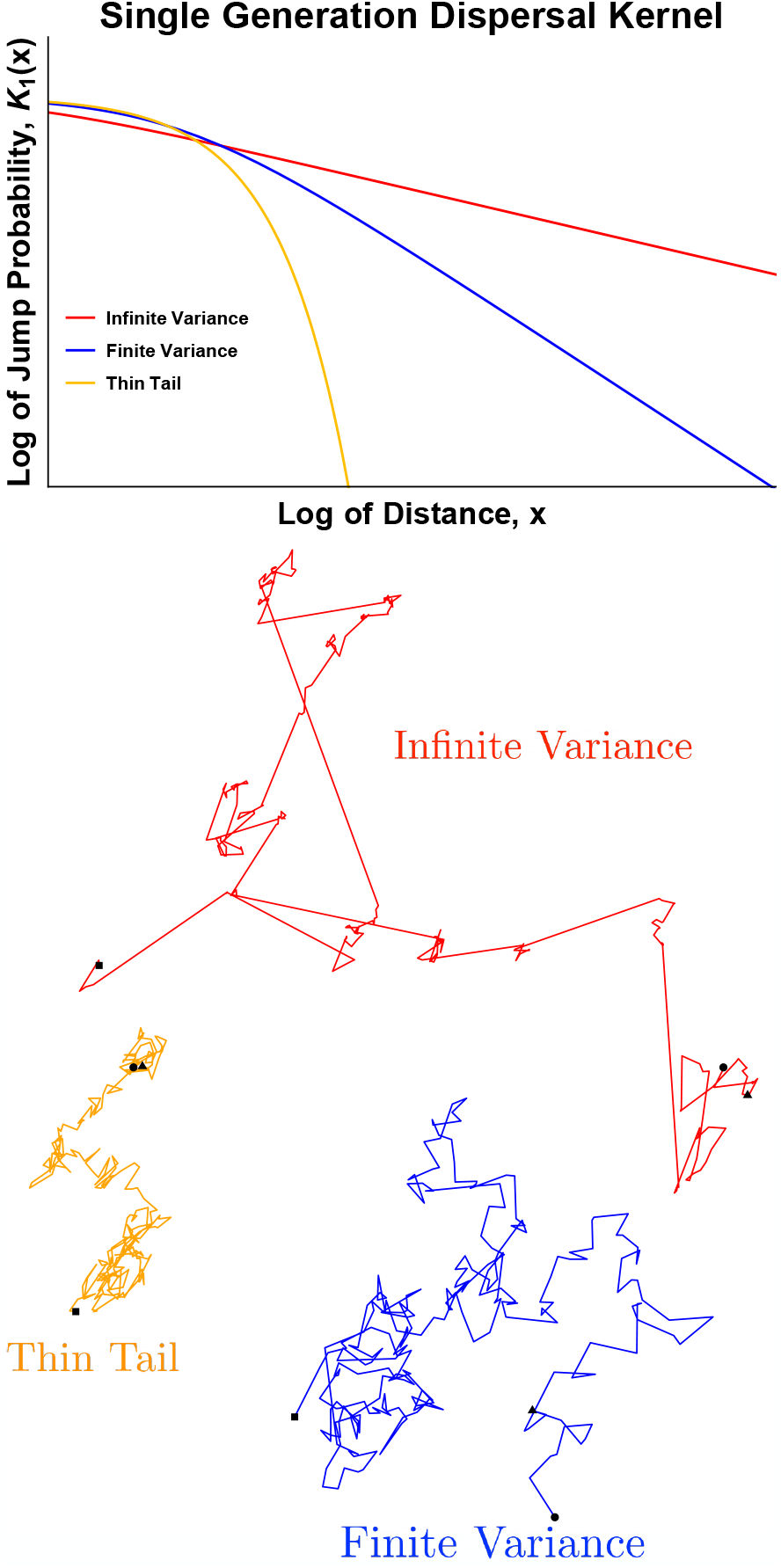
The tail of the dispersal distribution controls the size and number of long-range jumps. Top: Single-generation dispersal distributions. The orange curve shows a normal distribution, the classic thin-tailed distribution. The blue and red curves show distributions with power-law tails in which the probability of jumping farther than a long distance *y* is proportional to *y*^−*α*^. **Bottom: Two-dimensional random walks**. When the dispersal distribution is thin-tailed, the motion reduces to normal diffusion without any long-range jumps. When the dispersal distribution has a power-law tail, trajectories can jump large distances in a single time step. If the power-law tail is broad (*α* < 2), trajectories will have divergent mean squared displacement, and large jumps become noticeably more prevalent than for steep power laws with finite variance (*α* > 2). Circles mark the beginnings of the trajectories, triangles mark the positions after 10 jumps, and squares mark the ends. The generalized dispersal constants *D*_*α*_ are chosen such that the trajectories all have similar characteristic displacements at the time step marked by triangles. On shorter time scales, the diffusive trajectory tends to have the largest displacement, while on longer time scales the infinite variance trajectory in red tends to have the largest displacement.

The different lineage dynamics produced by power-law dispersal can have large effects on evolution, greatly accelerating both selective sweeps and range expansions, as in both cases individuals who make rare, long jumps into fresh patches can have very large numbers of descendants (Mollison 1972; Ibrahim *et al*. 1996; Mancinelli *et al*. 2003; Bialozyt *et al*. 2006; Brockmann and Hufnagel 2007; Wingen *et al*. 2007; Fayard *et al*. 2009; Ralph and Coop 2010; Hallatschek and Fisher 2014; Paulose *et al*. 2019; Paulose and Hallatschek 2020). This is true even for dispersal with *α* > 2 (Hallatschek and Fisher 2014). It is thus important to determine the tail of the dispersal distribution in natural populations. But for most populations, particularly non-animal ones, very little is known about dispersal, and the tail is particularly hard to observe directly (Koenig *et al*. 1996). We would therefore like to know if power-law dispersal leaves distinctive traces in patterns of isolation by distance from which it could potentially be inferred genetically (Nathan *et al*. 2003; Cayuela *et al*. 2018).

In this work, we explore the effects of power-law dispersal on isolation by distance in neutrally evolving, demographically stable populations with constant density. This problem was previously studied by Nagylaki (1976) in one dimension for moderately heavy-tailed dispersal with finite mean distance (*α* > 1), and Janakiraman (2017) studied an analogous problem in chemical physics and found complementary results. Nagylaki (1976) also considered Cauchy-distributed dispersal (*α* = 1) in one dimension, and Chave and Leigh Jr (2002) did the same for two dimensions. We unify and extend this work to cover arbitrary power-law dispersal tails in both one and two spatial dimensions and find simple asymptotic expressions for isolation by distance for both distant and nearby pairs of individuals. We also find how the distribution of the time to the most recent common ancestor of a pair of individuals depends on the distance between them. Our most important novel results are the expression (4) for relatedness between distant individuals under power-law dispersal with arbitrary *α*, which generalizes Nagylaki (1976) and Chave and Leigh Jr (2002)’s results beyond the special cases of *α* = 1 in *d* = 2 dimensions and 1 ≤ *α* < 2 in *d* = 1 dimension, and the expression (6) for relatedness between nearby individuals under broad power-law dispersal with *α* < *d*, which generalizes Chave and Leigh Jr (2002)’s result beyond the special case of *α* = 1 and *d* = 2. We also give heuristic derivations for the results in addition to precise mathematical ones, to make it easier to tell when they should hold in real biological populations that do not conform precisely to the mathematical models.

## Model

We consider two individuals sampled in the present a distance *x* apart, and trace their lineages backward through time until they coalesce. We assume that the distance that an individual moves in one generation is drawn from a distribution that falls off as a power law at long distances. For concreteness, we will mostly consider lineages that follow Lévy flights, a flexible, mathematically tractable way to model dispersal with power-law tails with *α* < 2 (Jespersen *et al*. 1999; Metzler and Klafter 2000; Metzler *et al*. 2009). Lévy flights occur naturally as the limit of any trajectory composed of many independent, identically distributed jumps, including the special case of diffusive motion in which the jumps have a finite variance. Apart from the special case of classic diffusion (for which the power-law tail disappears), these trajectories include rare long-range jumps and are governed by power-law kernels with infinite variance. We expect that most of our results are insensitive to the precise shape of the dispersal kernel and depend mostly on the tail behavior. Indeed, we see a good match between the asymptotic approximations and simulated populations dispersing according to three different kernels with the same tails but different short-range behavior: Lévy flights (for one-dimensional populations with *α* < 2), discrete approximations to F-distributions (populations with *α* > 2), and other discrete distributions with power-law tails (two-dimensional populations with *α* < 2); see Methods for simulation details.

In the main text, we will focus on the case of individuals living in a two-dimensional range; for the one-dimensional case, see the Methods section below. We will assume that the range size is large enough that it is effectively infinite; see the Discussion for more on this point. For *α* > 2, the motion of a typical lineage is approximately diffusive at long times. By this we mean that at time *t*, a typical lineage will have a displacement of 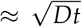. Here *D* is the diffusion constant, which is set by variance of the dispersal kernel. For *α* < 2, the dispersal kernel has infinite variance, and the typical displacement will instead be ≈ (*D*_*α*_*t*)^1/*α*^, which grows more quickly than the diffusive displacement at long times (Fig. 1). The parameter *D*_*α*_ is a generalized diffusion constant with units of distance^*α*^ /time, set by the typical scale (roughly, median) of the dispersal kernel. (For precise definitions of the Lévy flight dispersal kernel in terms of *α* and *D*_*α*_, see (17) and (47) for one and two dimensions, respectively.) For *α* < 2, the typical size of the maximum of *t* dispersal events is proportional to (*D*_*α*_*t*)^1/*α*^, and the probability that together they carry the lineage more than than a long dis-tance *y* ≫ (*D*_*α*_*t*)^1/*α*^ is proportional to *y*^−*α*^. For Lévy flights, *α* is known as the “stability parameter”. The stability parameter is identical to the tail exponent for *α* < 2, but in the limiting case *α* = 2, the Lévy flight reduces to ordinary diffusion, rather than a distribution with a power-law tail ∝ *y*^−2^. For the rest of this paper, when we refer to “*α* = 2”, we will mean diffusive motion. We will discuss dispersal kernels with power-law tails with exponent *α* > 2, but we will avoid dispersal kernels with exactly quadratic tails to minimize confusion.

When the two lineages encounter each other, they coalesce at rate proportional to 1/*ρ*, where *ρ* is the density of the population. Technically, in *d* = 2 dimensions, two lineages of infinitesimal size will never be at exactly the same position (Mörters and Peres 2010); the same is true for *α* ≤ 1 in *d* = 1 dimension (Bertoin (1996) p34). So really there must be some small distance *δ* within which lineages coalesce at a rate that is approximately 1/(*δ*^*d*^*ρ*), the inverse of the number of individuals within coalescence range. At these small scales, even the model of independent dispersal of lineages breaks down (Barton *et al*. 2010). But we will see below that this coalescence length scale does not affect isolation by distance on larger scales *x* ≫ *δ*. These considerations reflect the serious issues with finding coalescent models that correspond to forward-time models in continuous space (Felsenstein 1975). To avoid these issues, we treat our continuous model as being an approximation to an underlying discrete-space model of a square lattice of demes. It is this discrete model that we use in our two-dimensional simulations, while in one dimension we use the approximate continuous-space model; see the Simulation Methods below.

We are interested in the probability *ψ* of identity by descent of our sample pair as a function of the distance between them, *x*, which we will also refer to as the homozygosity or relatedness.

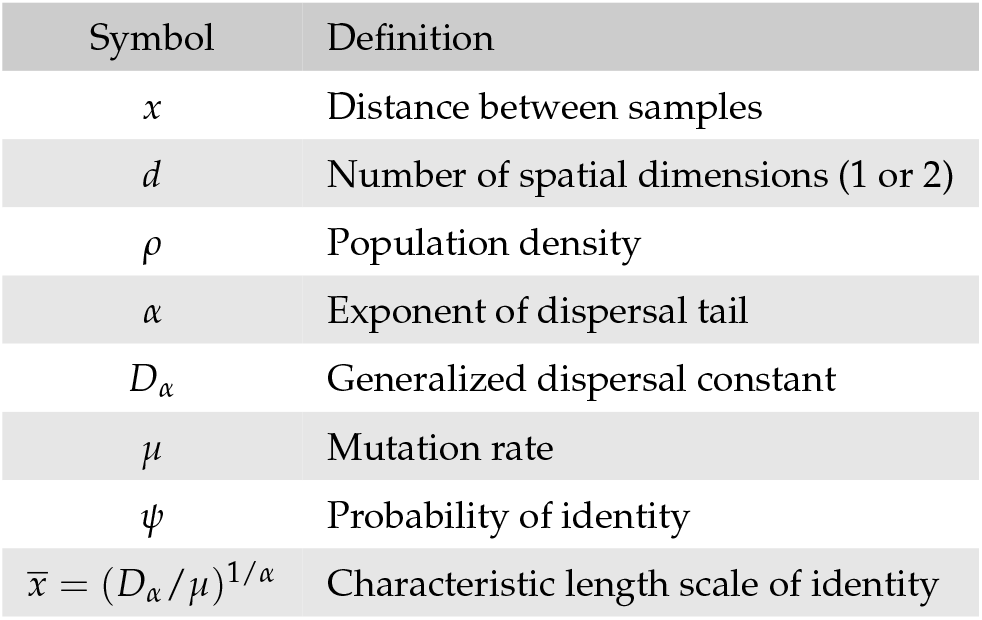

(See the Discussion for more on the interpretation of *ψ*.) If the time to their most recent common ancestor is *T* and the mutation rate *μ* is then *ψ* is given by:

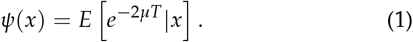

Although usually it is *ψ* rather than the coalescence time *T* itself that is directly observable, *T* is important for, e.g., determining whether it is reasonable to assume stable demography, so we will also find expressions for its probability density *p*(*t*|*x*).

## Results

In this section, we will describe our main results and provide brief sketches of the logic behind key features. We will focus on the scaling of identity *ψ* with the underlying parameters *α, D*_*α*_, *μ*, and *ρ*, mostly leaving numerical prefactors for the Methods. Roughly speaking, the basic intuition is that the sampled pair will be identical if their lineages coalesce within the past ∝ 1/*μ* generations. In this time, they will disperse a typical distance of order 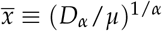, so this is the key length scale over which identity decays: pairs separated by 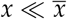 should be relatively closely related, while identity between pairs separated by 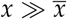 should be rare.

For the classic case of diffusive motion (*α* = 2), this length scale is 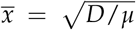, and the probability of identity falls off exponentially in one dimensional ranges (Barton *et al*. 2002):

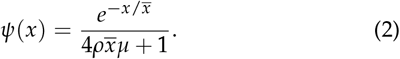

In two dimensions, it falls off falls off logarithmically for 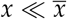 and exponentially for 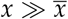 (Barton *et al*. 2002):

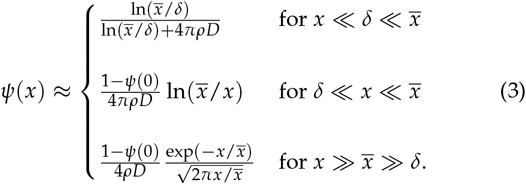

Here we generalize (2) and (3) to *α* ≠ 2, and find simple approximate expressions for *ψ* in different parameter regimes, illustrated in Fig. 2. At long distances, we find that *ψ*(*x*) has a universal form for all power-law dispersal kernels. Intuitively, power-law dispersal broadens the distribution of coalescence times for pairs at a given separation *x*, creating more overlap in the distributions for different *x* values (Fig. 3; see Appendix B for the equivalent schematic in one dimension).

**Figure 2.**
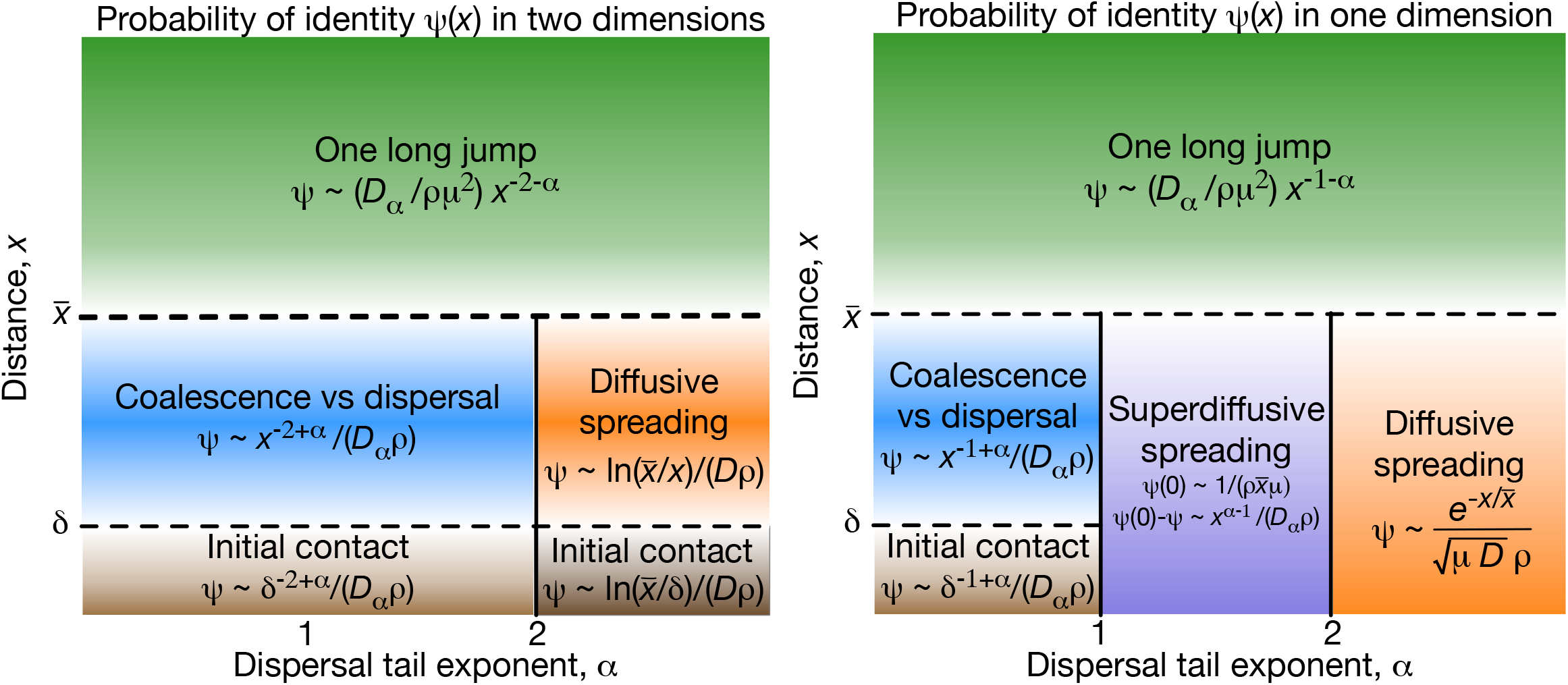
For power-law dispersal, the form of isolation by distance is universal at long distances. Approximate form for the probability of identity as a function of distance, *ψ*(*x*), for different dispersal kernel exponents *α*. Left panel shows results for *d* = 2 spatial dimensions, right panel shows *d* = 1 dimension. Different regimes of parameter space are labelled by their qualitative dynamics. We use “∼” to denote proportionality in the limit of large population density where *ψ*(0) ≪ 1. The key length scales are the characteristic length scale of identity, 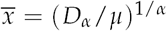, and the short distance *δ* at which coalescence can occur. Coalescence for distant pairs, 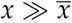, typically occurs via one long jump, which leads to the universal power-law scaling at large distances predicted by (4) (green). The form of isolation by distance among nearby pairs, 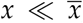, depends on *d* and *α*. For sufficiently heavy-tailed dispersal (*α* < *d*), nearby pairs typically either coalesce very quickly (at *t* ≪ 1/*μ*) or disperse far away from each other and mutate before coalescing, so the probability of identity is set by a competition between coalescence and dispersal and is nearly independent of the mutation rate ((6), blue). In this regime, identity still follows a power law with distance, although a shallower one than the power law at long distances, and with the opposite dependence on *α*. Finite-variance dispersal with *α* > 2 is effectively diffusive at short distances, with lineages typically reaching each other via many small jumps, so identity follows the classic diffusive predictions (2) or (3) (orange). In one dimension, there is an intermediate regime 1 < *α* < 2, in which nearby pairs typically reach each other by many small jumps but the non-diffusive nature of the jumps is still apparent ((10), purple). For *d* = 2 and *α* ≤ 1 in *d* = 1, the short-range details of coalescence become important for very close pairs ((3) or (7), brown). The solid vertical lines at *α* = 1 and 2 indicate sharp transitions, while the dashed horizontal lines at *x* = *δ* and *x* indicate smooth variation between asymptotic limits (see Figs. 4 and 5).

**Figure 3.**
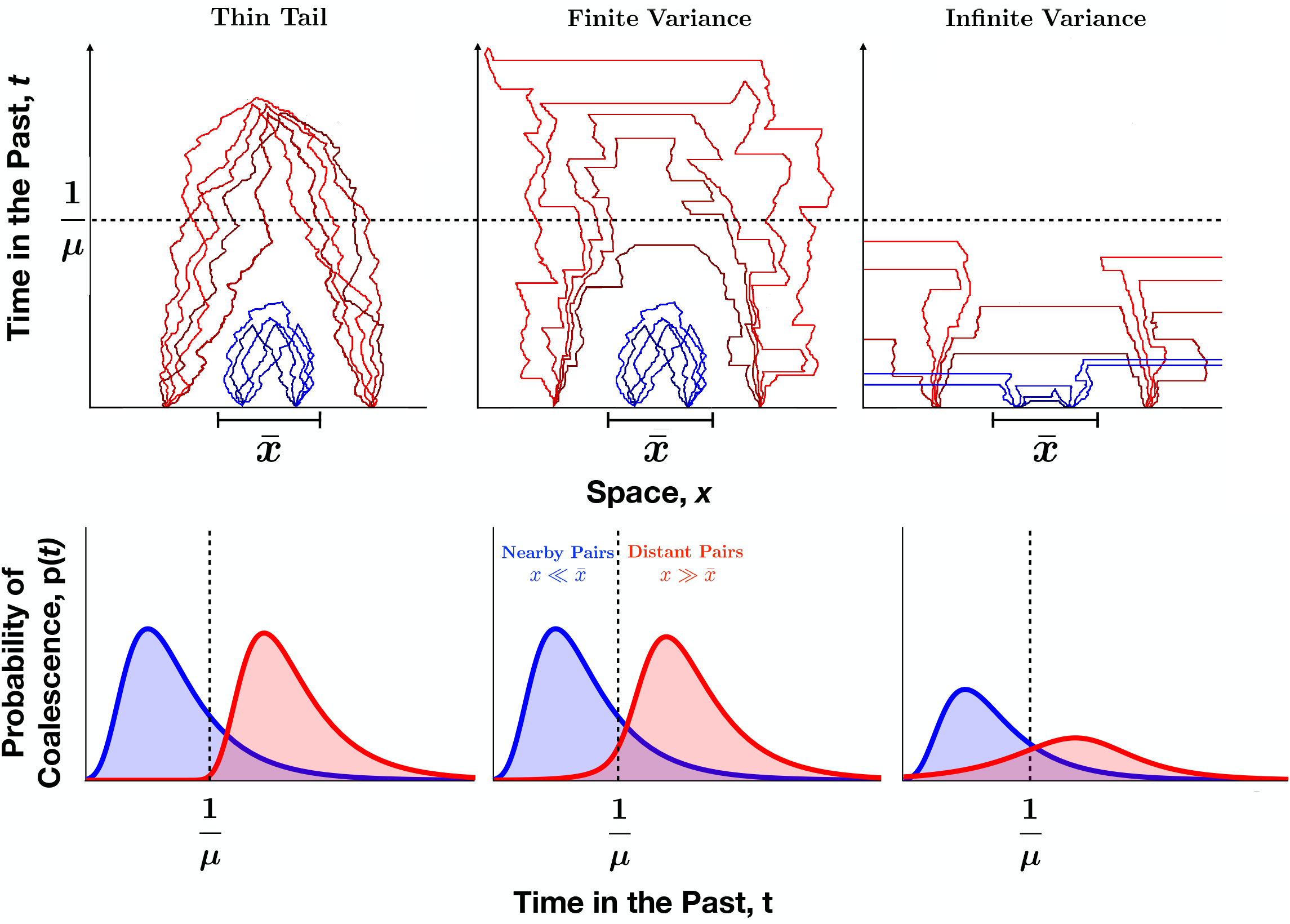
Long-range jumps affect when and where lineages coalesce. Qualitative illustrations of lineage dynamics and coalescence time distributions for each of the three dispersal regimes in two dimensions. Typical histories are shown for nearby samples (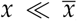, blue) and distant samples (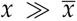, red). Left: For thin-tailed dispersal distributions (e.g., the normal distribution with *α* = 2), motion is effectively diffusive and separation *x* is a relatively good predictor of coalescence time. Center: For steep power-law dispersal distributions with finite variance (*α* > 2), large jumps broaden the spatial and temporal ranges over which lineages coalesce. Lineages at large separations 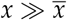 are occasionally able to coalesce at times comparable to 1/*μ*, while lineage dynamics at short distances are indistinguishable from thin-tailed dispersal. Right: For broad power-law dispersal distributions with infinite variance (*α* < 2), large jumps are common. This allows for the rapid coalescence of lineages at both small and large distances but also lets even nearby lineages jump very far away from each other and avoid coalescing until a much later time set by the range size (not shown).

### Distant pairs

For distant samples, 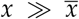, we expect substantial isolation by distance. For the pair to coalesce, their lineages must approach within *δ* of each other. The most likely way for this to happen before a mutation occurs is for one lineage to cover the distance in a long jump. In one dimension, the rate of such jumps is proportional to *D*_*α*_*x*^−*α*−1^ *δ*: a probability density proportional to *D*_*α*_*x*^−*α*−1^ for a jump to have size ≈ *x*, multiplied by a target zone of size *δ*. In two dimensions, the rate is ∝ *D*_*α*_*x*^−*α*−2^ *δ*^2^: there is an extra factor ∝ 1/*x* because the jump not only has to have the right distance but also the right direction, and now it must hit a target with area *δ*^2^. (Here we are focusing just on scaling behavior and neglecting 𝒪(1) numerical factors like 2 and *π*.) Combining the two expressions, the rate of jumps bringing the two lineages together is ∝ *D*_*α*_*x*^−*α*−*d*^*δ*^*d*^, where *d* = 1 or 2 is the spatial dimension. The probability that such a jump occurs in the time ∝ 1/*μ* before the lineages mutate is therefore approximately ∝ *D*_*α*_*x*^−*α*−*d*^*δ*^*d*^ /*μ*. The lineages must then coalesce within their neighborhood of ∝ *δ*^*d*^*ρ* individuals before they mutate. If mutation is frequent compared to coalescence (1/(*ρδ*^*d*^) ≫ *μ*), this occurs with probability ∝ 1/(*μρδ*_*d*_) ≪ 1. We therefore expect that the probability of identity is *ψ*(*x*) ∝ *D*_*α*_*x*^−*α*−*d*^*δ*^*d*^ /*μ*/(*μρδ*^*d*^) = *D*_*α*_*x*^−*α*−*d*^ /(*μ*^2^*ρ*), i.e., that there is a power-law dependence of identity on distance, with the same exponent as the dispersal probability density. We can generalize this expression slightly to also include the case in which mutation is rare compared to local coalescence by including a correction factor that depends on the probability of identity *ψ*(0) for a co-located pair:

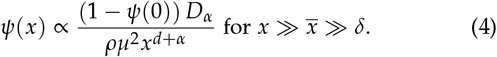

See the Methods for a derivation of (4), including the omitted constant of proportionality, which depends on the details of the dispersal distribution. For *d* = 1 and 1 ≤ *α* < 2, (4) was derived by Nagylaki (1976) (his Eq. (37)); our result extends this to two dimensions and arbitrary *α*. We confirm (4) with simulations and numerical analysis (see Fig. 4 for two dimensions and Fig. 5 for one dimension). As shown in Fig. 4 and Fig. 5, even steep power-law kernels with *α* > 2 and finite variance lead to a matching power law in *ψ*(*x*). (4) tells us that the probability of identity between widely separated individuals is far higher than would be predicted under diffusive dispersal ((2) or (3)), and that identity falls off more slowly at long distances for dispersal distributions with fatter tails (smaller *α*).

**Figure 4.**
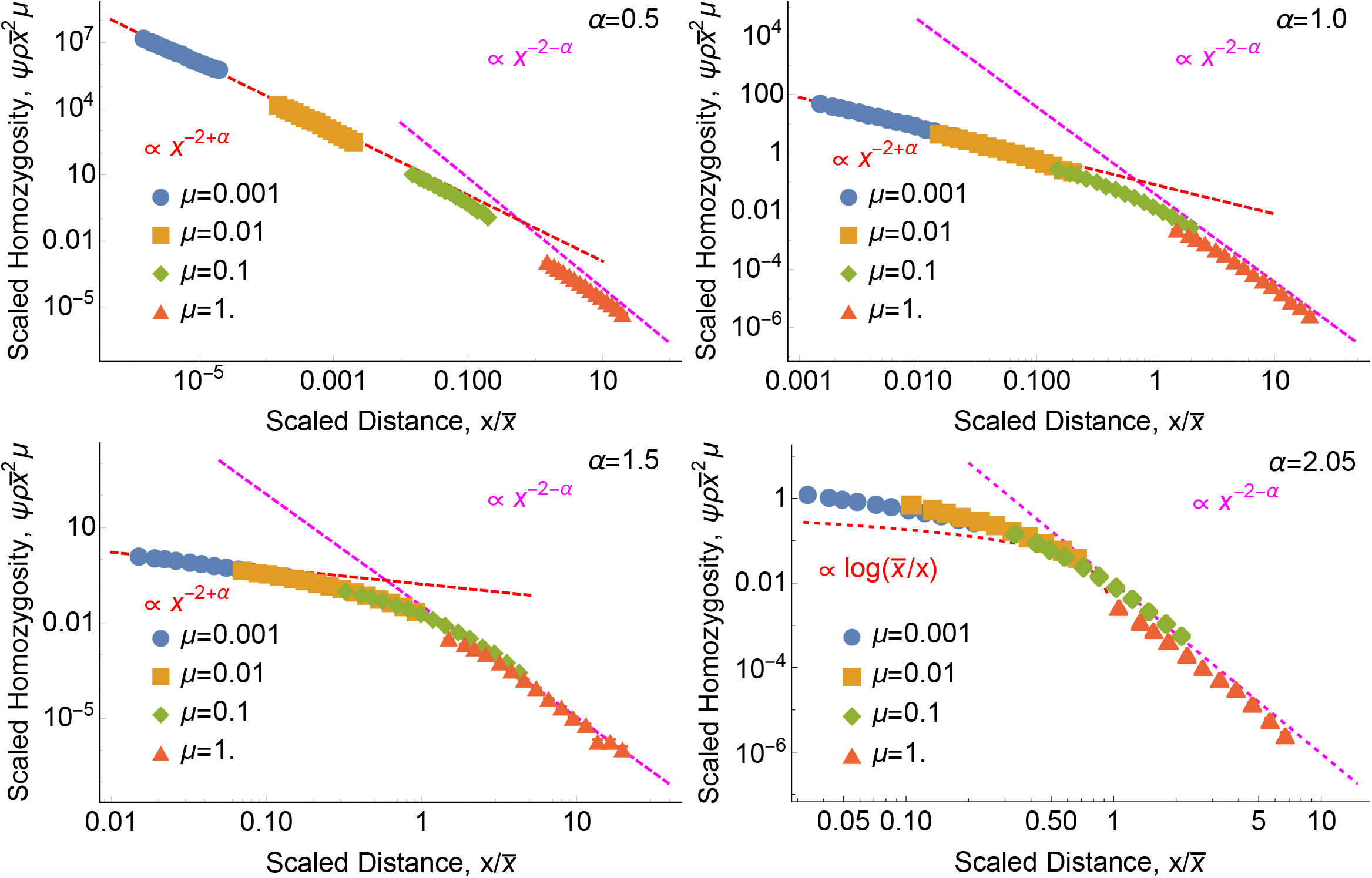
Isolation by distance in two dimensions has the same power-law tail as dispersal. Each panel shows the scaled probability of identity between a sampled pair of individuals, 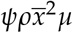, as a function of the scaled distance 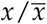 between them. Points show discrete-space simulation results and magenta lines show the power law that emerges at large distances (4) (see (62) for prefactors). For *α* < 2, red curves show the short-range power-law behavior predicted by (6). For *α* = 2.05, the red curve shows the classic diffusive prediction, (3). Notice that even for this finite-variance case that looks diffusive on short scales, the underlying non-diffusive power law is apparent at long scales. *ρ* = 1 in all panels. For these parameter values, identity is rare even for co-located individuals, *ψ*(0) ≪ 1. In this and all following figures, error bars (which in this figure are smaller than the points) show 68% percentile bootstrap confidence intervals (see Methods).

**Figure 5.**
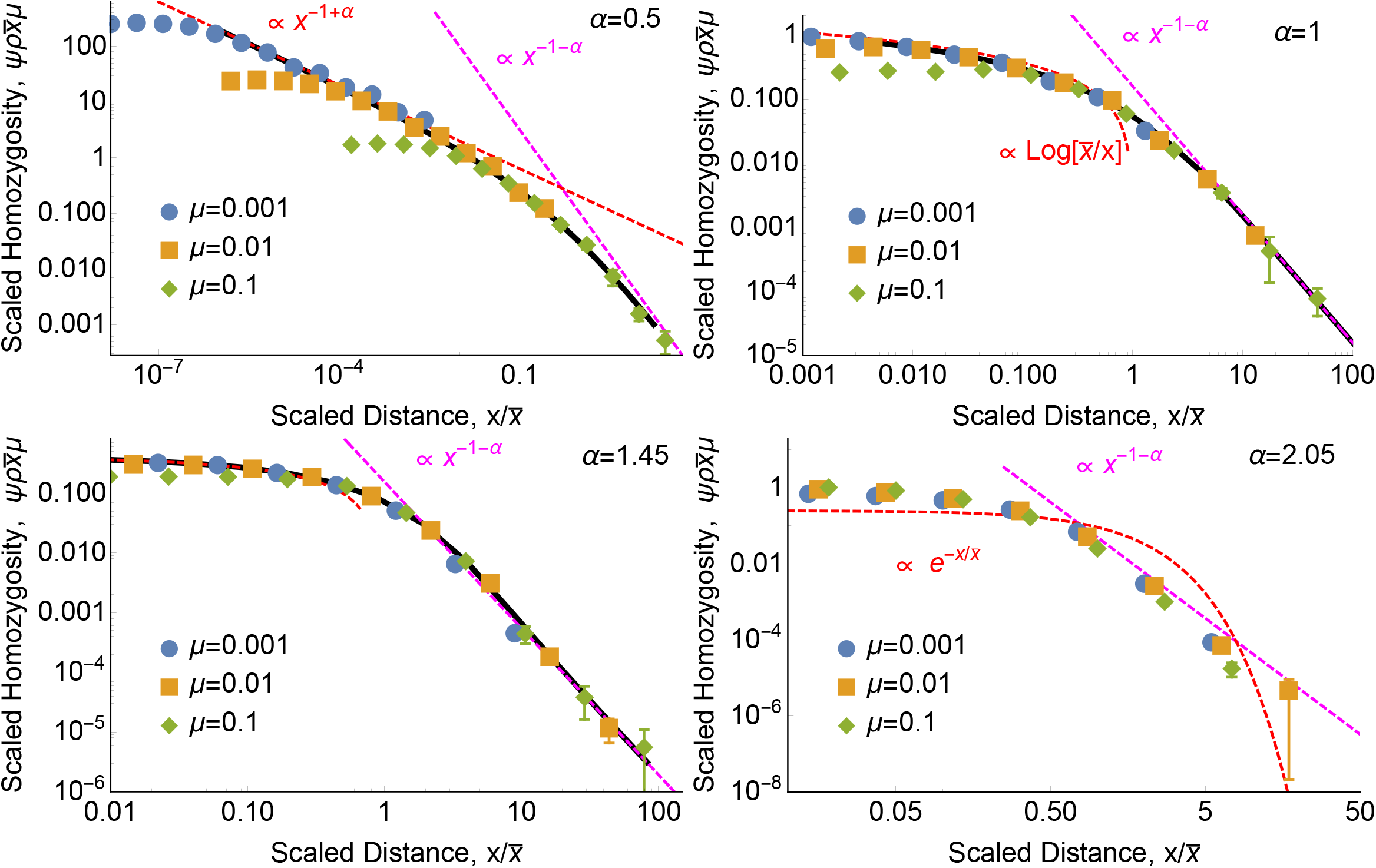
Isolation by distance in one dimension also has the same power-law tail as dispersal. As in Fig. 4, each panel shows the scaled probability of identity between a sampled pair of individuals, 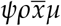, as a function of the scaled distance 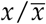 between them. Points show simulation results and magenta lines show the same long-distance power law (4). Black curves show numerical solutions of *ψ*(*x*) calculated from (33) with *δ* = 0 and 1 − *ψ*(0) → 1. (As in Fig. 4, parameters are chosen such that *ψ*(0) ≪ 1.) Red curves show the asymptotic behavior predicted at short distances by (6) for *α* = 0.5, by (8) for *α* = 1, by (9) and (10) for *α* = 1.45, and by the classic diffusive result (2) for *α* = 2.05. Again, the finite-variance case looks diffusive at short distances but follows the underlying power law of dispersal at long distances. *ρ* = 100 in all panels, and data with *ρ* = 10 and *ρ* = 1 (not shown) yield indistinguishable plots.

### Nearby pairs

The lineages of nearby pairs 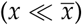 do not need to take a long jump to find each other before mutating. In fact, the long-range jumps in the tail of the dispersal kernel can actually carry the pair farther away from each other and make it harder to coalesce. Intuitively, the pair of lineages take time ∝ *x*_*α*_ /*D*_*α*_ to disperse across the distance between them. (Here, we are assuming *α* ≤ 2; steep power laws with *α* > 2 will be approximately described by the diffusive *α* = 2.) From that time on, they are roughly uniformly likely to be anywhere within a range of expanding radius ∝ (*D*_*α*_*t*)^1/*α*^. They thus coalesce at a decreasing rate ∝ (*D*_*α*_*t*)^−*d*/*α*^ /*ρ*. To find the probability that they coalesce before mutating, we integrate this rate over time, starting from the time ∝ *x*_*α*_ /*D*_*α*_ when they first approach each other, out to the time ∝ 1/*μ* by which they mutate:

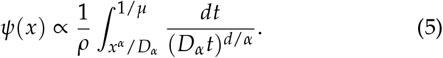

This describes the regime where identity is rare, *ψ*(*x*) ≪ 1, before it saturates at 1 for sufficiently low density *ρ*. Notice that the initial separation *x* enters into (5) only through the lower limit of integration, while the mutation rate *μ* enters only through the upper limit. This implies that the derivative *ψ* ′(*x*), the rate at which identity decreases with distance at short distances 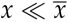, should be approximately independent of *μ*. The behavior of (5) depends strongly on the comparison between the tail exponent *α* and the dimension *d*, and so we will treat the different cases separately below.

### Nearby pairs: broad power-law kernels

We will first consider broad power-law tails, with *α* < *d*. In this case, the exponent *d*/*α* of the denominator in (5) is > 1, and the integral is therefore dominated by the region near the lower limit of integration (short times). In other words, nearby lineages are likely to either coalesce very quickly or to disperse across the whole range before coalescing (Bertoin (1996) p34, Palyulin *et al*. (2014)). This “now-or-never” dynamic has the interesting effect of making the local probability of identity by descent independent of the mutation rate, because the main competition is between coalescence and heavy-tailed dispersal rather than between coalescence and mutation. Explicitly, we obtain *ψ*(*x*) ∝ 1/(*ρD*_*α*_*x*^*d*−*α*^), with the fact that the upper limit of integration is 1/*μ* rather than infinity only negligibly decreasing *ψ*(*x*). So as with distant pairs, *ψ*(*x*) follows a power law, although a different one from the long-distance 1/*x*^*d*+*α*^. We calculate *ψ*(*x*) more carefully in the Methods to find that it is given by:

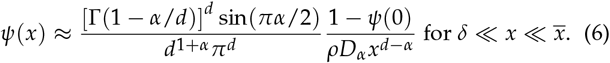

We include the complicated numerical prefactor for completeness, but it will typically be of order one, and given that the model is an idealization of any real system and that the effective density *ρ* will only be known very roughly, we do not expect it to be very important in practice. Note that when *d* = 2 and *α* = 1, (6) reduces to Eq. (A6) in Chave and Leigh Jr (2002). We confirm (6) with simulations (Fig. 4 and Fig. 5).

There are two key features distinguishing (6) from the classic thin-tailed results. First, the power-law behavior indicates that *ψ* can change rapidly even at short distances 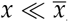, unlike the exponential or logarithmic behavior predicted by (2) and (3), respectively. Second, as mentioned above, *ψ* is approximately independent of the mutation rate *μ*. We consider the interpretation of this point in the Discussion below. There is also an interesting contrast between (6) and (4): *α* appears with opposite signs in the two exponents. Thus, while broadening the tail of the dispersal distribution (by decreasing *α*) makes isolation by distance weaker for distant pairs (i.e., the dependence of *ψ* on *x* becomes weaker in (4)), it makes isolation by distance stronger for nearby pairs. This is because distant pairs rely on long-range jumps to bring them together, while for nearby pairs such jumps are likely to push them apart and prevent coalescence.

The power law in (6) makes it diverge at very short distances, where it breaks down. Instead, for individuals within the same deme, *x* < *δ, ψ*(*x*) flattens out, with *ψ*(*x*) approaching *ψ*(0). Roughly speaking, individuals coalesce at rate 1/(*ρδ*^*d*^) and disperse outside of coalescence range at rate of about *D*_*α*_*δ*^−*α*^. When coalescence is faster, probability of identity is high, *ψ*(0) ≈ 1, while when dispersal is faster it is low, *ψ*(0) ≈ 1/(*ρδ*^*d*^)/(*D*_*α*_*δ*^−*α*^) = 1/(*ρD*_*α*_*δ*^*d*−*α*^). A more careful calculation gives (see Methods):

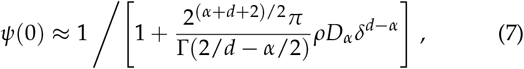

although these numerical factors depend on the details of the coalescence kernel. We confirm (7) with simulations for *d* = 1 (Fig. 6).

**Figure 6.**
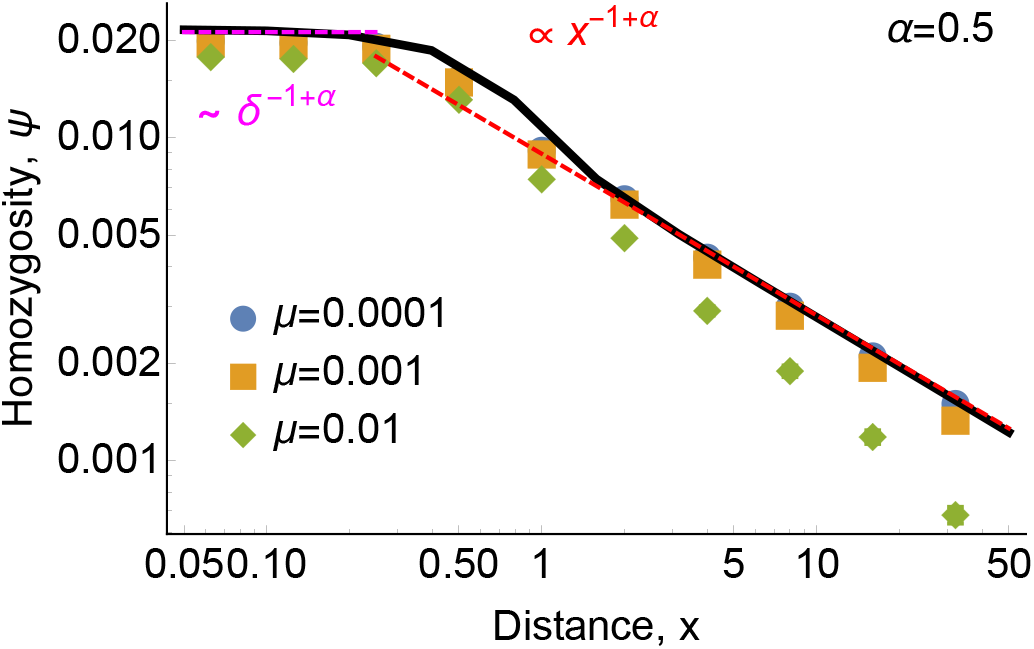
For heavy-tailed dispersal with *α < d*, relatedness at short distances is independent of mutation rate but is sensitive to the length scale *δ* of coalescence at very short distances. Nearby lineages at 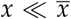 either coalesce quickly and are identical, or jump very far away from each other and never coalesce. Points show continuous-space simulation results in one dimension, and red and magenta lines show the asymptotic predictions of (6) and (41), respectively. The black curve shows a numerical solution of *ψ*(*x*) calculated from (33) with *μ* = 10^−4^. *ρ* = 100 in all plots, and data with *ρ* = 10 and *ρ* = 1 (not shown) yield indistinguishable plots.

In populations with discrete generations, (6) and (7) both break down if *x* is small enough that the individuals have a substantial probability of coalescing in a single generation. More precisely, the expression *x*^*α*^ /*D*_*α*_ in the lower limit of integration in (5) assumes that *x* is large enough to be in the tail of the single-generation dispersal kernel. See “Breakdown of models at small scales” in the Methods below for a discussion of alternative expressions when this is not satisfied.

### Nearby pairs: marginal kernels

The marginal case *α* = *d* is actually the most familiar, as it includes the classic case of diffusion in two dimensions. For *α* = *d* = 2, evaluating (5) recovers (3), up to numerical constants. As mentioned above, finite-variance dispersal kernels with *α* > 2 will effectively be described by *α* = 2, so this case actually describes a broad region of parameter space. This is shown in Fig. 4 for *α* = 2.05.

There is also the one-dimensional marginal case, *α* = *d* = 1. In the Methods, we show that like the classic marginal case *α* = *d* = 2, it produces logarithmic isolation by distance among nearby pairs:

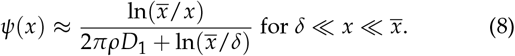

We confirm (8) with simulations (Fig. 5). We believe that (8) and the associated expression for co-located pairs (43) are novel, but that their relevance to natural populations is limited, as they apply only in one dimension and for *α* exactly one. On the other hand, for selective sweeps such marginal exponents can actually describe the dynamics in a substantial region of parameter space when there is not a clean separation of scales (Hallatschek and Fisher 2014), and it is possible that something similar could be true here, in this case perhaps when *α* is close to one and *x* is not too large compared to *δ*.

### Nearby pairs: moderate power-law kernels in one dimension

In two dimensions, the two cases listed above are the only possibilities. For one-dimensional habitats *d* = 1, there is an additional regime *α* > 1. For finite-variance dispersal *α* ≤ 2, this reduces to the classic diffusive case described by (2). For moderately broad power laws 1 < *α* < 2, the integral in (5) is dominated by the region near the upper limit of integration (long times). This gives a leading term 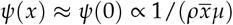 that is independent of *x*. The spatial dependence enters as a correction from the finite lower limit of integration, *ψ*(0) − *ψ*(*x*) ∝ *x*^*α*−1^/(*ρD*_*α*_). A more detailed analysis (see Methods) recovers the result of Nagylaki (1976) (Eq. (39)) for *ψ*(0):

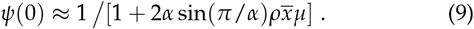

The leading distance dependence was found by Janakiraman (2017) (Eq. (C1)) for an analogous problem in chemical physics:

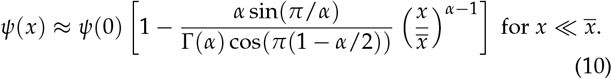

Notice that this matches the scaling predicted by the intuitive argument. We confirm (9) and (10) with simulations and numerical analysis (Fig. 5).

While (10) is similar to the pattern for finite-variance dispersal (2) in that *ψ* is only slowly changing for small *x*, the details are slightly different, in that (10) predicts a linear correction to *ψ* with *x* to leading order, while (10) predicts a leading term ∝ *x*^*α*−1^. But this is only a modest difference and it is unclear if it could be detected in data from natural populations; the long-distance behavior discussed above presents a much clearer contrast. Because the constants in (10) are the result of behavior over many jumps, allowing the lineage displacements to approach a Lévy stable distribution, they should be independent of the details of the dispersal kernel. However, they are typically of order one and are unlikely to be practically important.

## Discussion

### Summary and related work

Limited dispersal produces a correlation between spatial and genetic distance (Wright 1946; Malécot 1975; Slatkin 1991; Slatkin and Arter 1991; Slatkin 1993). While most previous models have only considered diffusive dispersal, dispersal can be heavy-tailed in many natural populations. Nagylaki (1976) was the first to generalize classic diffusive models of isolation by distance by allowing dispersal distance in one-dimensional populations to have a power-law tail (with 1 ≤ *α* < 2). This groundbreaking work has largely been neglected, likely because of the lack of data to which to compare it at the time; it was last cited by Chave and Leigh Jr (2002), who extended the results to two dimensions for the special case of Cauchy flights (*α* = 1) in a paper modeling ecological diversity. Recent studies suggest that heavy-tailed dispersal may in fact be common (Willson 1993; Clark 1998; Atkinson *et al*. 2002; Baguette 2003; Brockmann *et al*. 2006; Dai *et al*. 2007; Fric and Konvicka 2007; Devaux *et al*. 2007; Aguillon *et al*. 2017; Vallaeys *et al*. 2017), and we hope that this paper will reintroduce these classic results to population genetics now that the field may have sufficient data to apply them. We also extend this previous work to all tail exponents *α* in both one and two dimensions. We find that, for all *α*, power-law dispersal leads to much more heavy-tailed relatedness than diffusive dispersal, with relatedness having the same power-law tail in distance as the dispersal kernel. This is true even for steep kernels with finite variance. In this case, even though a diffusive approximation can fit the pattern of isolation by distance between nearby individuals, it will greatly underestimate the degree of relatedness between distant individuals.

In addition to the power-law tail of relatedness at long distances, we also find that heavy-tailed dispersal with *α* < *d* (where *d* is the number of spatial dimensions) produces a different, shallower power-law decay of relatedness at short distances. This is in contrast to the pattern predicted under diffusive models, in which relatedness depends only weakly on distance until falling off exponentially at longer distances ((2) and (3)). The patterns predicted by heavy-tailed dispersal seems more consistent with examples from natural populations (e.g., Aguillon *et al*. (2017)) in which there is clear isolation by distance even at short distances, but still substantial relatedness at long distances.

Barton *et al*. (2013b) and more recently Forien (2022) and Forien and Wiederhold (2022) have considered a similar model for populations evolving according to a spatial Λ-Fleming-Viot process in which large extinction-recolonization events causing dispersal and coalescence of many lineages have a power-law distribution of spatial extents. Our model differs from Barton *et al*. (2013b) and Forien (2022) in that dispersal of different lineages and coalescence are all independent, and in that coalescence is purely short-range. Very recently, Forien and Wiederhold (2022) have constructed an alternative spatial Λ-Fleming-Viot model that can accommodate a closer match to our analysis, as well as other patterns of dispersal and coalescence. Specifically, they have two different distributions for the locations of offspring and parents, allowing dispersal and coalescence to be partially decoupled. They demonstrate that our scaling results can apply both to their model and to the original (Forien 2022). An even closer match between models could be achieved by having two classes of events: dispersal would be primarily driven by very mild but long-range events affecting lineages with a small probability decaying as a power law from the epicenter of the event, while coalescence would primarily be driven by intense but localized events. Note that while this would give an exact continuous-space backwards-time model to which our analysis would apply, the spatial Λ-Fleming-Viot only has an approximate forwards-time counterpart (Barton *et al*. 2013b), while the discrete-space simulations we use correspond exactly to a forwards-time model.

### Possible application in dispersal inference

Our results connect power-law dispersal kernels and the resulting patterns of pairwise relatedness. Standard methods for inferring dispersal from pairwise measures of relatedness or autocorrelations in allele frequency typically assume either thin-tailed, diffusive motion (Rousset 1997, 2000; Robledo-Arnuncio and Rousset 2010; Ringbauer *et al*. 2017; Bradburd *et al*. 2018) (perhaps with recent long-range admixture (Bradburd *et al*. 2016)) or a small number of discrete demes (Slatkin 1991; Whitlock and McCauley 1999; Rousset and Leblois 2011; Petkova *et al*. 2016; Al-Asadi *et al*. 2019; Lundgren and Ralph 2019). Methods using cline theory to infer dispersal from the width of hybrid zones make similar assumptions about the motion of lineages being diffusive (Barton 1983; Barton and Hewitt 1985; Sotka and Palumbi 2006; Rieux *et al*. 2013; Gagnaire *et al*. 2015; Cayuela *et al*. 2018). These methods can be adjusted to accommodate occasional infinite-range dispersal by treating it analogously to mutation (Rousset 2007), but this does not give rise to the kinds of new functional forms for isolation by distance found here. Methods for non-stable demographies based on historical biogeography or coalescent theory tend to also assume a small number of discrete demes (Sanmartín *et al*. 2008; Ree and Smith 2008; Hey 2010).

Other genetic methods such as parentage analysis are better equipped to infer heavy-tailed dispersal on continuous ranges, but these techniques require exhaustive sampling of the population to ensure that the parents of each individual can be located (Adams *et al*. 1992; Jones and Ardren 2003; Bacles and Ennos 2008; Wang and Santure 2009). More recent methods for pollen dispersal have been developed that allow for the inference of heavy-tailed dispersal without the need for exhaustive sampling, but knowledge of maternal genotypes for all sampled individuals is still required (Austerlitz *et al*. 2004; Robledo-Arnuncio *et al*. 2006). For plant species where this data is available, our results could serve as the basis for complementary inference methods. While the pollen dispersal methods are focused on inferring the dispersal kernel over a single generation, isolation by distance reflects the history of dispersal over many generations, so a comparison of the results could reveal changes in dispersal over time. For species where no such pedigree data is available, continuous-space inference methods based on the model developed here could allow for the presence (or absence) of heavy-tailed dispersal to be inferred for the first time.

One key open question is to what extent it is possible to detect the genetic traces of rare heavy-tailed dispersal in natural populations, and if so how well the form of heavy-tailed dispersal (e.g., the tail exponent *α*) can be determined. Austerlitz *et al*. (2004) were able to detect heavy-tailed pollen dispersal in *Sorbus torminalis* tree populations using parentage analysis and the seed-specific TwoGener method. There was substantial uncertainty in their estimates of *α*, but this was based on data from 2000, and sequencing data has grown enormously in the two decades since. The combination of vastly more data and a complementary inference method based on our results could allow for reasonable precision in estimating *α*. However, estimating power laws is difficult in general (Clauset *et al*. 2009), and it seems unlikely that one would be able to confidently distinguish between, say, *α* = 1.1 and *α* = 1.2; on the other hand, this distinction is not likely to be very important in real populations with finite range sizes, while the difference between *α* ≈ 1.2 and thin-tailed dispersal is likely to be quite important and should be apparent in data.

### Scaling “mutation” by varying the length of genomic blocks

Along with predicting characteristic scaling of identity by descent with distance, our results predict characteristic scaling with the mutation rate *μ*, and also a scaling of the typical length scale of identity *x* with *μ*. While mutation rate cannot be varied directly as distance can, *μ* here should be understood as referring to the mutation rate in a block of non-recombining genome, and so a wide range of effective *μ* values can be scanned by considering identity by descent in blocks of varying size (Weissman and Hallatschek 2017). This will be valid as long as recombination is rare relative to mutation, or even if recombination is frequent as long as “*μ*” is understood to mean the sum of the block mutation and recombination rates, and recombination events can be reliably detected using marker loci. In this case, *ψ* would be the probability that a region of fixed length is identical by descent, as opposed to being interrupted by either mutation or a recombination “junction” (Fisher 1954). (And if one has a population sequenced only at marker loci, *ψ* could be defined as just the probability that the region is free of junctions, neglecting mutation, in which case “*μ*” would be just the block recombination rate.) This suggests that it should be possible to measure identity by descent statistics corresponding to *μ* values ranging over five orders of magnitude in a single sample (Harris and Nielsen 2013). A natural extension of the present theory would be to consider the expected *distribution* of lengths of segments of identity by descent as a function of distance (Ralph and Coop 2013; Ringbauer *et al*. 2017) under different dispersal kernels.

This ability to increase the effective *μ* is crucial to dispersal inference in general. In natural populations, per-nucleotide pairwise genetic diversity is low (*π* ≪ 1, (Buffalo 2021)), i.e., *ψ*(*x*) ≈ 1 even for large *x* when we using the single-site mutation rate for *μ*. This means that finite range effects play a strong role on these time scales, limitations in dispersal have a correspondingly small effect, and there is little hope of detecting the scaling behavior we find here. But increasing the effective *μ* by looking at longer genomic regions restricts our time horizon for coalescence. This makes *ψ* insensitive to the right tail or even the bulk of the coalescence time distribution, and instead focuses on unusually rapid coalescence events, where isolation by distance is much more apparent (Ralph and Coop 2013; Ringbauer *et al*. 2017; Aguillon *et al*. 2017). Ideally, one can find an optimal range of genomic lengths, short enough that identity by descent is fairly frequent among nearby individuals but long enough that it is much less frequent among distant individuals.

Seen in this light, our result that the identity *ψ* for nearby pairs of individuals is nearly independent of *μ* for *α* < *d* ((6) and (42)) means that we predict that any blocks of identity by descent between nearby individuals should be very long. They should not extend over the whole genome, however, because our definition of “nearby”, 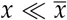, contains an implicit *μ* dependence through 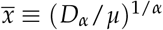. Solving the condition 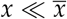 for *μ*, this suggests that the size of blocks of identity by descent between a pair of individuals a distance *x* apart should be broadly distributed up to a genomic length proportional to *D*_*α*_ /*x*^*α*^, beyond which mutation and recombination become effective in breaking them up.

As noted above, (5) implies that the rate at which identity decreases with distance at short distances, *ψ*′ (*x*) for 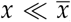, is independent of *μ* for all *α* and *d* as long as *ψ*(0) ≪ 1. (This can also be seen in the full results by differentiating (2), the middle line of (3), (6), and (10).) This suggests that if the probability of identity as a function of distance were plotted for blocks of genomic length *l*, the curves for different values of *l* (i.e., different values of “*μ*”) should be approximately parallel at short distances. Because this pattern does not depend on the details of the dispersal kernel, deviations from it would indicate that other processes, e.g., demographic fluctuations or local adaptation, were important in determining isolation by distance.

### Effect of finite range size

Our analysis has neglected effects due to finite range size, although our two-dimensional simulations take place in habitats of finite length. To better understand the effect of finite range size, we can consider a pair of individuals sampled from random locations within a habitat of length *L*; the mean coalescence time between them would then be the “effective population size”, *N*_*e*_. The mean pairwise genetic diversity *π* is directly proportional to this time, *π* = 2*N*_*e*_*μ*. The pair will typically be sampled a distance of about *L* from each other, and so it will typically take a time of order *L*_*α*_ /*D*_*α*_ for their lineages to overlap in space. At this point the ancestry is effectively well-mixed, and coalescence takes time proportional to the total population size *N* = *L*^*d*^*ρ*, where *d* = 1 or 2 is the dimension of the habitat. For *L*^*α*^ /*D*_*α*_ ≪ *L*^*d*^*ρ*, the mixing time has little effect, while for *L*^*α*^ /*D*_*α*_ ≳ *L*^*d*^*ρ*, the mean coalescence time is substantially higher than one would expect in the panmictic limit, *N*_*e*_ > *N*. For thin-tailed dispersal, *α* = 2, structure substantially increases the mean coalescence time in a one-dimensional habitat of length *L* ≳ *Dρ* (Maruyama 1971), while in two dimensions the effect of space depends only on the local neighborhood size *Dρ* (Maruyama 1972), up to logarithmic corrections (Cox and Durrett 2002). The effect of population structure thus either increases with the spatial extent of the population (at fixed density) or is nearly insensitive to it. With heavy-tailed dispersal, however, we see a new qualitative pattern. For *α* < *d*, i.e., for broad power laws, the effect of structure on mean time to coalescence counterintuitively becomes *weaker* as the range size *L* grows, because *L*^*α*^ /*D*_*α*_ grows more slowly than *L*^*d*^*ρ*.

This new pattern suggests that the structure of genetic variation in space might be quite different for *α* < *d* than it is for the classic *α* = 2 models. In the classic diffusive models, if spatial structure is weak enough that the effective population size is not much more than the census size, *N*_*e*_ ≈ *N*, then even nearby lineages must typically wander over the whole range before coalescing. Conversely, if nearby individuals are frequently identical by descent (by which we mean coalescing in time ≪ *N*_*e*_), then structure must be strong enough to make *N*_*e*_ ≫ *N*, an unusual situation in natural populations. For *α* < *d*, on the other hand, movement can be slow relative to coalescence on short scales, allowing for substantial local identity by descent, while still being rapid on long scales and keeping *N*_*e*_ ≈ *N*.

### Outlook

Our use of stable distributions for the dispersal kernel has been partly motivated by the fact that any isotropic single-generation dispersal kernel will converge to a stable one if it is repeated over many independent generations. But as we have noted, this is only true asymptotically, and in any real population there will be correlations across generations, spatial inhomogeneities, shifts in dispersal over time, limits due to finite range size, and many other effects that cannot be captured by a stable distribution. It is therefore better to see it as a simple reference model, possibly one step closer to reality than the purely diffusive one, that can serve as a background against which to measure all these other effects. Whether it is worth adding the extra parameter *α* to the simple diffusive model with *α* = 2 will depend in part on whether the population of interest has a range large enough compared to typical single-generation dispersal distances for the tail of the dispersal distribution to matter.

What other processes could produce similar patterns to heavy-tailed dispersal? One obvious one is if individuals are performing something more like a “Lévy walk” than a Lèvy flight, in which dispersal in any one generation is thin-tailed but can be correlated across many generations (Zaburdaev *et al*. 2015). Such an effect can be produced at the level of alleles by hitchhiking on beneficial substitutions (Allman and Weissman 2018). But this should be readily distinguishable from neutral heavy-tailed dispersal by considering the distribution of relatedness across multiple individuals and loci—hitchhiking will produce heavy-tailed relatedness at the same few loci across all individuals, whereas neutral effects will be more evenly distributed. It is an open question whether other neutral processes, in particular demographic fluctuations, might produce similar patterns. We believe that the increase in sequence data from natural populations means that the time has come to develop theories which seriously connect spatial patterns of genetic diversity and the full range of plausible underlying dynamics, beyond the standard simplest cases.

## Methods

### Simulation methods in two dimensions

All simulation code and displayed data are available at https://github.com/weissmanlab/Long_Range_Dispersal. We simulate our model in two stages. First, for each value of present-day separation *x*, dispersal constant *D*_*α*_, and tail parameter *α*, we simulate dispersal of the lineages, ignoring coalescence and mutation. Then, for each value of *ρ* and *μ*, we calculate the expected homozygosity and coalescence time distribution for each simulated trajectory. We then average over many independent trajectories. This two-part method has two advantages. First, the same dispersal simulations can be used for calculating the homozygosity and coalescence time distribution for multiple choices of *ρ* and *μ*. Second, the latter part of the method, in which conditional expectations are calculated for previously generated paths, is entirely deterministic, reducing computational cost and noise. Note that this backward-time model corresponds exactly to a forward-time model of Wright-Fisher reproduction within demes and dispersal among demes according to the simulated kernel; this exact correspondence is only possible because we are using a discrete-space model.

We simulate lineage motion using a discrete time random walk,

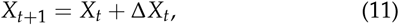

where *X*_*t*_ represents the position of a lineage at a given time (ignoring coalescence., i.e., assuming *ρ* → ∞), and the displacement, Δ*X*_*t*_, is a vector of integer-valued random variables drawn from the dispersal distribution at each integer time *t*. We use the GNU Scientific Library’s efficient pseudorandom generators for both stable distributions and the F-distribution (Galassi *et al*. 2009). Because these are available only for the one-dimensional distributions, we draw radial distances using the one-dimensional distributions and then select a direction in which to move uniformly at random.

To draw increments Δ*X*_*t*_ for *α* < 2, we first draw a positive random number from the continuous one-dimensional Lévy alpha-stable distribution:

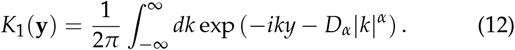

We then convert this to a two-dimensional vector y by drawing a direction uniformly at random. Finally, to keep lineages on a discrete lattice of demes, we round y to the nearest pair of integers, i.e., the closest point in ℤ^2^, to obtain Δ*X*_*t*_. In the GNU Scientific Library, the scale of the Lévy alpha-stable distribution is parameterized not by *D*_*α*_ but rather by *c* ≡ (*D*_*α*_)^1/*α*^, the characteristic spread *c* of each lineage after one generation (*t* = 1). We use *c* = 10 for all two-dimensional simulations with *α* < 2. Notice that (12) does not match the two-dimensional Lèvy alpha-stable distribution ((47) below) that we use for our analytical approximations, so the match between the analytical results and the simulations shows that the results are robust to the details of the dispersal kernel.

To simulate steeper tails with *α* > 2, we follow the same procedure as in the previous paragraph, but instead of drawing the initial continuous dispersal distance *y* from (12), we draw it from a degenerate F-distribution:

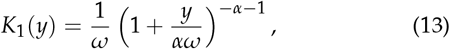

where *ω* scales the characteristic single-generation dispersal distance. (Technically, we obtain *y* by drawing from the GNU Scientific Library F-distribution with degrees of freedom *ν*_1_ = 2 and *ν*_2_ = 2*α* and scaling the result by *ω*.) Because this kernel has finite variance, the central limit theorem implies that at long times the bulk of the displacement distribution approaches that of a diffusive kernel, with dispersal constant *D* given by one-quarter the single-generation variance in position:

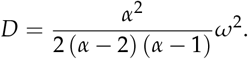

We use *D* = 200 for all two-dimensional simulations with *α* > 2.

For each pair of simulated dispersal trajectories {*x*_*t*_}, we then compute the path-specific distribution of coalescence times *p*({*x*_*t*′≤*t*_}), i.e., the probability of coalescing at and not before time *t*, and the path-specific mean homozygosity *ψ*({*x*_*t*′ ≤ ∞_}), i.e., the probability that lineages following these exact trajectories have not mutated before coalescence:

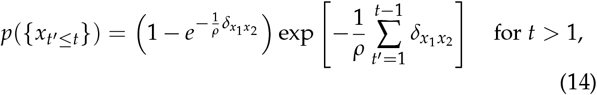

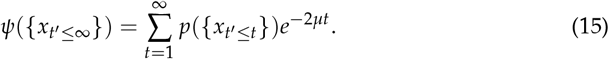

We start (14) and (15) at *t* = 1 because we assume that the individuals are sampled immediately after dispersal, so no coalescence takes place at 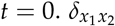 in (14) is the Kronecker delta function:

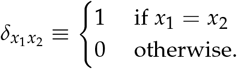

For every time-step the lineages spend in the same deme, there is a probability of coalescence 1 − exp(− 1/*ρ*). Note that this means that we interpret *ρ* as a local effective population size, rather than as the actual census size, e.g., *ρ* = 1 is interpreted as coalescence at unit rate, rather than certain coalescence in one generation. We do not expect that changing *ρ* to be the census size would significantly change our results. Note also that because we calculate only pairwise quantities, any value of *ρ* can be matched exactly in a forward-time model with deme census size *N* > *ρ* by increasing the variance in offspring number appropriately.

We then average (14) and (15) across all simulated trajectories with present-day separation *x* to obtain *p*(*t*|*x*) and *ψ*(*x*). All error bars in plots show 68% confidence intervals, as determined by the percentile bootstrap with 1000 bootstrap samples (Davison and Hinkley 1997). At large distances, the distribution of the probability of identity across sample trajectories is highly skewed, with most trajectories having very low probabilities of identity, but a few having the lineages rapidly jump close to each other and having a high probability of identity. This means that we cannot quantify the uncertainty in our estimates using, for example, the standard error of the mean, but it also means that we must simulate many independent trajectories to get good enough coverage for the bootstrap to be accurate (Chernick 2011).

For the simulations of mean homozygosity *ψ* shown in Fig. 4, we simulate 10^7^ independent runs of 1000 generations each for each combination of present-day separation *x* and tail parameter *α*. We also apply periodic boundary conditions, with the range size extending from − 5000 to 5000 along both dimensions of the discrete lattice. Our choice of range size is significantly larger than the maximum value used for present-day separation between pairs (*x* = 237). For the largest mutation rate considered, *μ* = 1, coalescence before mutation is extremely rare, and so we increase the number of independent runs to 10^8^ (with the number of generations reduced to ten).

### Simulation methods in one dimension

Our one-dimensional simulation methods are very similar to those used in two dimensions, with only three significant differences that together make them much simpler. First, instead of having to track two random walks, one for each lineage, we can just track the difference between their positions. Secondly, we take space to be continuous rather than discrete, i.e., we do not round the difference in positions and do not require the difference to be exactly zero for coalescence. Note that, as mentioned in the Model section above, this means that these simulations are only meant as approximations to some underlying model that has an exact forward-time counterpart. Finally, we take the range size to be effectively infinite, limited only by the maximum size of double-precision numbers.

Explicitly, we again simulate lineage motion using a discrete time random walk,

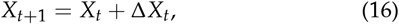

where *X*_*t*_ now represents the signed distance between two lineages at a given time, and the step size, Δ*X*_*t*_, is a real valued random variable drawn from the dispersal distribution at each integer time *t*. For *α* < 2, we use Lévy alpha-stable distributions for dispersal, with each lineage’s single-step displacement having probability density *K*_1_(*y*) given by:

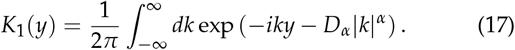

The quantity we actually simulate, Δ*X*_*t*_, the increment in the signed distance between the two individuals, is the sum of the two lineages’ independent jumps. Thus its probability density *K* is the convolution of *K*_1_ with itself:

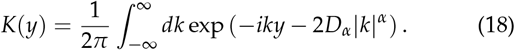

Because the distributions are stable, *K* differs from *K*_1_ only by an extra factor of two in the dispersal constant.

To simulate steeper tails with *α* > 2, we again use a degenerate F-distribution, drawing Δ*X*_*t*_ from the two-sided distribution:

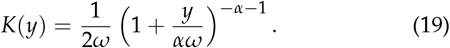

At long times, the bulk of the displacement distribution approaches that of a diffusive kernel, with dispersal constant *D* equal to half the mean squared single-generation displacement of one lineage:

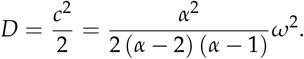

For each simulated trajectory {*x*_*t*_}, we then compute the path-specific distribution of coalescence times *p*({*x*_*t* ′ ≤*t*_}) and path-specific mean homozygosity *ψ*({*x*_*t* ′ ≤∞_ }):

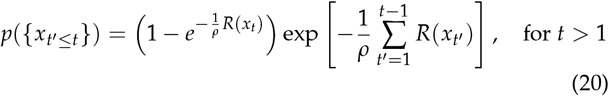

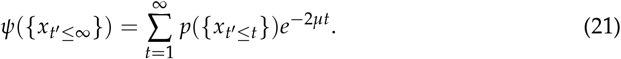

*R*(*x*) in (20) is a rectangular function representing a uniform rate of coalescence of all lineages within a distance *δ*:

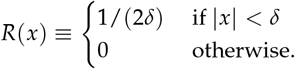

For every time-step the lineages spend in this region, there is a probability of coalescence 1 − exp [− 1/(2*ρδ*)]. We discuss issues with the microscopic interpretation of this model after we introduce our analytical model below.

Unconditioned values *p*(*t* | *x*) and *ψ*(*x*) are again obtained by averaging across all simulated trajectories. Error bars in plots show 68% confidence intervals, as determined by the percentile bootstrap with 10^4^ bootstrap samples.

We set *δ* = 0.5 for all simulations in one dimension. For the simulations of mean homozygosity *ψ* shown in Fig. 5, we simulate 250,000 independent runs of 1000 generations each for each combination of present-day separation *x* and tail parameter *α*. We set the dispersal constant *D*_*α*_ indirectly by setting the characteristic spread *c* of two lineages after one generation (*t* = 1), *c* = (2*D*_*α*_)^1/*α*^, to be fixed at *c* = 250 for *α* < 2, and *c* = 179.68 for *α* = 2.05. For the largest present-day separations, *x* = *e*^10^ and *e*^11^, coalescence within 1000 generations is very rare, so we increase the number of runs to 1.25 × 10^6^. For Fig. 6 and Fig. 8, we choose *D*_*α*_ such that *c* = 0.2, and simulate 10,000 independent runs of length 1000 generations each.

For the simulations of the cumulative distribution of coalescence times *P*(*t*) shown in Fig. 7, we set present-day separation *x* = 0 and generate 10,000 independent trajectories of 1.5 million generations each for each combination of *c* and tail parameter *α*. We set *c* = 3.59 for *α* > 2, *c* = 5 for 1 < *α* < 2, and *c* = 1 for *α* ≤ 1.

**Figure 7.**
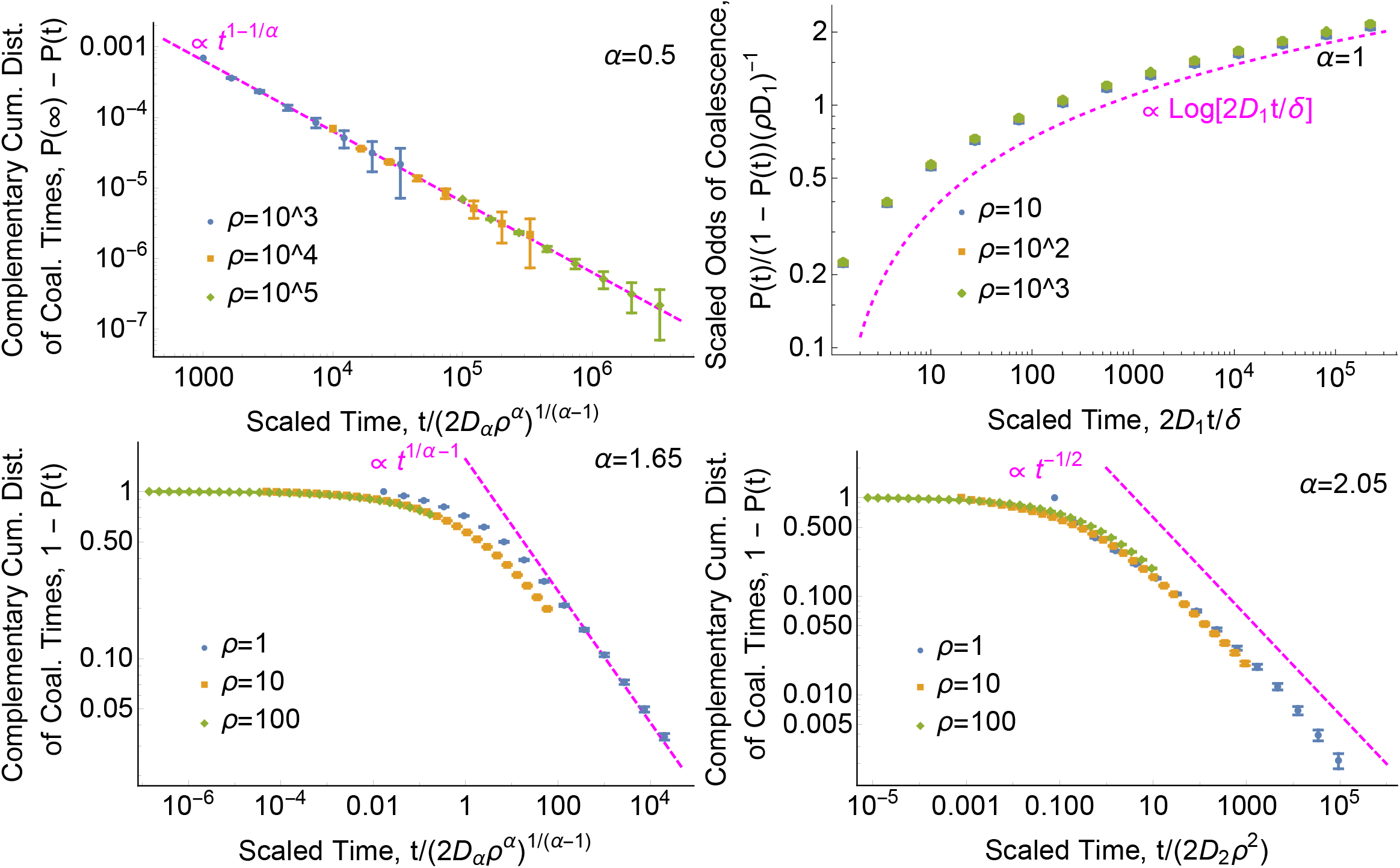
The distribution of coalescence times has a power-law tail. Points show one-dimensional simulation results. Dashed magenta curves show the asymptotic predictions (in order of increasing *α*) (67), (71), (68), and (70). Time is scaled to dimensionless units. See Simulation Methods section for *D*_*α*_ values. We show statistics based on the cumulative distribution *P*(*t*) rather than the density *p*(*t*) because simulation estimates for the latter are very noisy. **Top left: for *α <* 1 in one dimension, the distribution of coalescence times is proportional to the probability of lineages being nearby, *K*(0|*t*) ∝ *t*^1−1/*α*^**. Plot shows *P*(∞) − *P*(*t*) rather than 1 − *P*(*t*) because lineages can disperse infinitely far away from each other and avoid coalescing entirely, i.e., *P*(∞) < 1. We use the simulated value of *P*(*t* = 10^6^) to approximate *P*(∞). This empirical value deviates from the continuous-time prediction (41) by ≈ 30% due to differences in the amount of coalescence in the first few generations (see “Breakdown of models at small scales”). **Top right: the distribution of coalescence times has a logarithmic tail for *α* = 1 in one dimension**. In this marginal case, lineages do eventually coalesce even in infinite ranges, but can take extremely long to do so. **Bottom left: for 1 *< α <* 2, the distribution of coalescence times in one dimension decays more quickly than the probability of lineages being nearby**. The coalescence time probability density has a power-law tail, *p*(*t* | *x*) ∝ *t*^1/*α*−2^. This deviation from the scaling of the dispersal kernel at long times is due to the high probability of previous coalescence events. **Bottom right: for *α >* 2, the coalescence time distribution may approach the diffusive limit**. The scaling of 1 − *P* appears to be close to that of the diffusive prediction, (70), but there is at least a difference in prefactor, perhaps again due to different probabilities of coalescence at very recent times. Present-day separation *x* was set to zero for all simulation results shown.

### Analytical model in one dimension

#### Generic dispersal

We want to find a tractable analytical approximation to the model described above. For recurrent motion, the lineages will sometimes be in exactly the same place, and we can model coalescence with a *δ* distribution, i.e., as taking place at rate 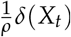. For transient motion, however, they will never coincide (Palyulin *et al*. 2014), and we must allow coalescence to take place at a finite distance. Let the coalescence kernel be some probability density 𝒩 (*x*) symmetric about *x* = 0 and with width ∼ *δ*, with coalescence taking place at rate 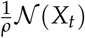. The *δ*-distribution is just the limit of 𝒩 as *δ* goes to 0, so we can treat the two cases together. Forien (2022) avoids this issue by using a spatial Λ-Fleming-Viot model in which dispersal and coalescence are produced by the same heavy-tailed process, but this leads to dispersal distances for an individual’s offspring being strongly correlated rather than independent (Barton *et al*. 2013b). Very recently, Forien and Wiederhold (2022) have examined a spatial Λ-Fleming-Viot model that addresses this by having separate dispersal and coalescence kernels, allowing a mathematically rigorous treatment in continuous space. We instead interpret our continuous-space model as an approximation to an underlying discrete-space model as in our two-dimensional simulations. As mentioned in the Simulation Model section, this means that the continuous-space model must break down at some point, which we discuss below in “Breakdown of models at small scales”.

The the probability density of coalescence times for lineages with initial displacement *x* is then:

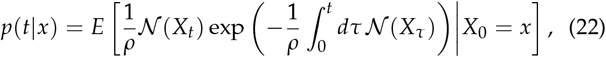

and the probability of identity is its Laplace transform:

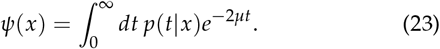

There are several different ways to derive an explicit expression for *ψ* from (23), including balancing mutation, coalescence, and dispersal over an infinitesimal time step (Malécot 1975; Barton *et al*. 2002) or, for Lévy flights, using a fractional diffusion equation (Janakiraman 2017) (see Appendix A). Here we start with a generalization of Barton and Wilson (1995)’s expression for *p*(*t* | *x*) that is valid for any two-lineage dispersal kernel *K*, which is defined as the convolution of *K*_1_ with itself. Assuming that 𝒩 (*x*) = *δ*(*x*):

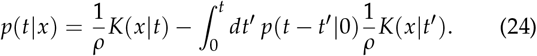

To interpret (24), notice that the first term is the probability of coalescing at time *t* neglecting the possibility that the lineages have coalesced more recently. The second term corrects for these more recent coalescences: for every trajectory where the lineages coincide at *t*′ < *t*, we subtract off the probability that the lineages would coalesce at *t*′ and then again exactly at *t*. Notice that we do not need to correct again for lineages that coincide three times, at *t*′′ < *t*′ < *t*: the factor of *p* guarantees that each trajectory is weighted appropriately.

We can immediately find a simple expression for *ψ* for recurrent dispersal by taking the Laplace transform of (24):

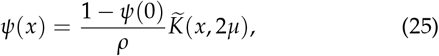

where tilde denotes the Laplace transform. Plugging in *x* = 0, we can solve (25) for *ψ*(0) and express *ψ*(*x*) purely in terms of the dispersal kernel:

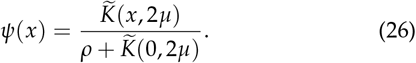

Note that, for diffusive dispersal, (25) reduces to the classical Wright-Malécot formula for isolation by distance (Barton *et al*. 2002).

For transient dispersal, we must consider a coalescence kernel of finite width, and (24) generalizes to:

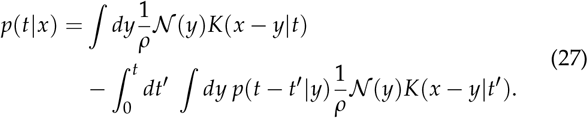

(27) is exactly the same as (24) except that now we must integrate over possible locations *y* of coalescence at both *t* and *t*′. Taking the Laplace transform of (27) now gives:

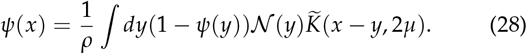

To simplify (28), we can make the approximation that 1 *ψ*(*y*) is nearly constant over all separations |*y*| ≲ *δ* where 𝒩 (*y*) is non-negligible, allowing us to pull it out of the integral:

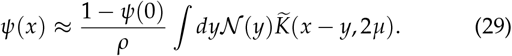

This approximation will necessarily be accurate when identity is low, *ψ*(0) ≪ 1 because 1 − *ψ* will be close to 1 for all *y*. However, for 1 − *ψ*(0) ≪ 1, the approximation can become inaccurate; we discuss this below. At long distances 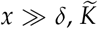 will also be roughly constant in the integral, and we simply recover (25), although now only as an approximation:

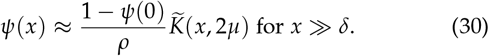

We see that the details of the short-range behavior only affect the long-range probability of identity by descent through the overall factor 1 − *ψ*(0) (Barton *et al*. 2002). Mathematically, the main challenge is to find simple expressions for *ψ*(0) and especially 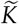.

Because (30) is invalid for *x* = 0, we cannot solve it directly for *ψ*(0) as we could with (25), and so we must also work with (29). We can simplify the convolution in (29) by taking the spatial Fourier transform ℱ{·}:

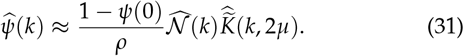

where 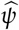 and 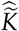 are the Fourier transforms of *ψ* and 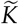.

#### Lévy flight dispersal

For Lévy flights, the characteristic function is 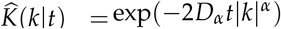 and the Fourier-Laplace transform is 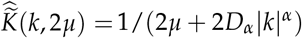. (31) for 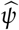 is correspondingly simple:

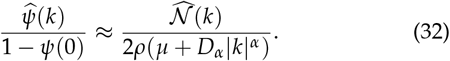

To get an explicit expression for *ψ*, we need to specify a form for the coalescence kernel 𝒩. We will use a normal distribution with standard deviation *δ*, which has the simple Fourier trans-form 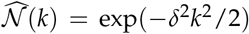. Then we can invert the Fourier transform in (32):

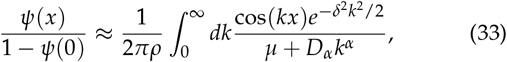

which can be re-expressed in dimensionless units as

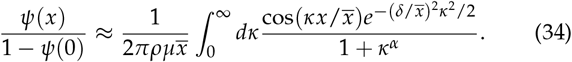

Examining (33), we see that the power-law tail in the integrand can be cut off either when oscillations in the cosine factor become rapid at *k* ∼ 1/*x* or by the normal factor at *k* ∼ 1/*δ*. As long as we are sampling pairs that are outside the immediate range of coalescence, *x* ≫ *δ*, the former cutoff will happen at lower *k*, and therefore the normal factor can be neglected (by setting *δ* = 0), leaving (in dimensionless form):

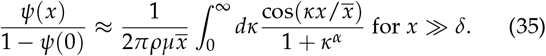

(35) can equivalently be derived directly from (30) by substituting in the Lévy flight dispersal kernel and writing 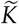 as the inverse Fourier transform of 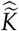.

We can solve (34) for *ψ*(*x*) by first evaluating it at *x* = 0 to find *ψ*(0); we do this below. But it is interesting that the ratio Ψ(*x*) ≡ *ψ*(*x*)/(1 − *ψ*(0)) has the simplest relationship to the underlying parameters, as shown by Rousset (1997) for short-range dispersal. Ψ is closely related to Rousset (2000)’s statistic *a*_*r*_ : *a*_*r*_ = Ψ(0) − Ψ(*r*). It is also related to the expected pairwise *F*_ST_ between demes separated by *x*:

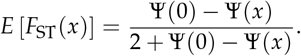

### Probability of identity for distant pairs 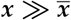, α < 2

For large *x* ≫ (*D*_*α*_*t*)^1/*α*^, the dispersal kernel has a simple asymptotic form for *α* < 2 (Nolan (2018), Theorem 1.12):

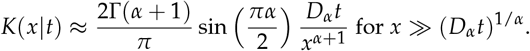

Plugging this into (30) and evaluating the Laplace transform gives the probability of identity for distant pairs, which was originally found by Nagylaki (1976):

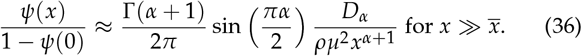

### Probability of identity for distant pairs 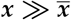, α > 2

There is no stable distribution with *α* > 2, but in discrete-time models such as the one we use in our simulations, we can consider single-generation jump kernels *K*(*y*|1) with power-law tails with *α* > 2. These will approach a diffusion with diffusion constant *D* = Var(*K*)/4. At long distances 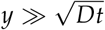, however, the tail will still be dominated by the probability of taking a single large jump (Vezzani *et al*. 2019), so for 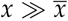, we will have *K*(*x*| *t* ≲ 1/*μ*) ≈ *K*(*x*|1)*t*. Plugging this into (30) and evaluating the Laplace transform gives:

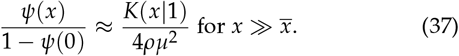

For the F-distribution kernel (19) used in the simulations, this is

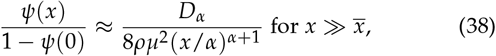

where we have defined *D*_*α*_ *ω*^*α*^ /generation, i.e., *D*_*α*_ has the same value as *ω*^*α*^, but its dimensions are now length^*α*^ /time. (38) is confirmed by simulations (Fig. 5, *α* = 2.05). We can then use the classic diffusive expression for *ψ*(0) to get an explicit expression for probability of identity at large distances:

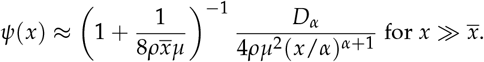

### Moderately heavy-tailed dispersal, 1 < α < 2

For *α* > 1, (25) and (35) are exact for all *x* (when *δ* = 0). Evaluating (35) for *x* = 0 gives *ψ*(0):

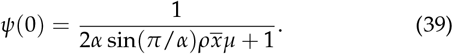

Plugging (39) into (36) lets us solve for *ψ*(*x*) at large distances 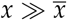:

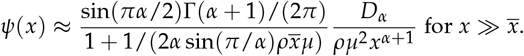

For 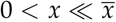, Janakiraman (2017) (Eq. (C1)) found that to leading order *ψ* falls off as:

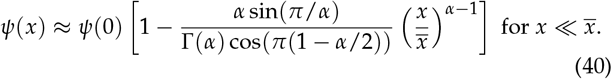

When *α* = 2, the above expression is equivalent to the classic diffusive result for 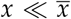, which can be found by integrating (28) with *δ* = 0:

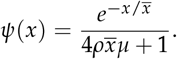

### Very heavy-tailed dispersal, α < 1

For *α* < 1, the finite width *δ* of the coalescence kernel is important for determining *ψ*(0). Setting *x* = 0 in (34) gives:

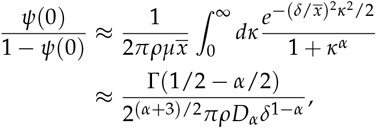

where in evaluating the integral we have assumed that 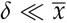, i.e., that the mutation rate is not extremely large. We see that on small scales, the probability of identity by descent is independent of the mutation rate (Fig. 6), i.e., there is a large probability that individuals from the same deme are differentiated even for very low mutation rates:

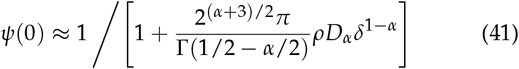

Very heavy-tailed dispersal of nearby lineages causes them to quickly wander away from each other, and for infinite range size many pairs will never coalesce. While (41) is only accurate for *ψ*(0) ≪ 1, the independence from mutation rate should persist even for large *ψ*(0).

Plugging (41) for *ψ*(0) into (36) gives an explicit expression for the probability of identity of distant pairs:

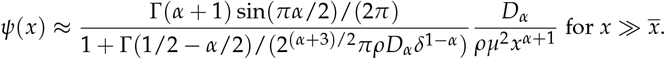

For pairs that are nearby but still well outside of coalescence range, 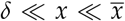, the integral in (35) is dominated by *κ* ≫ 1 and is approximately:

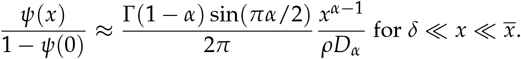

Again, the probability of identity is independent of the mutation rate to lowest order. Substituting in (41) gives an explicit expression for *ψ*:

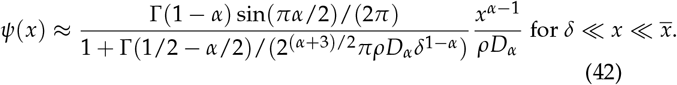

While *ψ*(*x*) is independent of *μ* only for *α* < 1, note that the rate at which *ψ*(*x*) changes for small *x*, 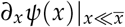, is independent of *μ* for all *α* when *ρ* is large.

### Marginal case α = 1

The analysis of the marginal case *α* = 1 is essentially the same as for *α* < 1 above, but we have separated it out because the form of the final expressions is very different. As with *α* < 1, the finite coalescence width *δ* is important for *x* = 0:

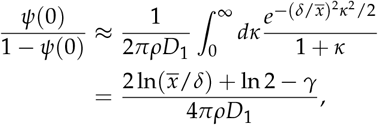

where *γ* ≈ 0.58 is Euler’s constant. Again assuming 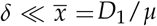, the constant terms in the numerator can be neglected and *ψ* is approximately:

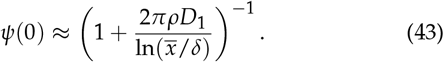

Recall that the approximation we used to derive (43) ((29)) is only justified when *ψ*(0) 1.

For pairs that are nearby but still well outside of coalescence range, 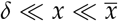, (35) gives:

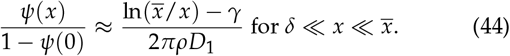

Plugging the expression (43) for *ψ*(0) into (44) and (36) gives explicit expressions for *ψ*(*x*) at both short and long distances:

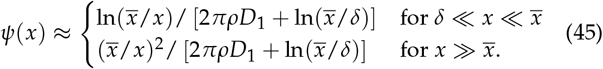

### Analytical model in two dimensions

#### Generic dispersal

For generic dispersal, the solution for *ψ* in two dimensions can again be found from (28), now with the integral over two spatial dimensions. The Fourier transform 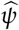 has the same form as the one-dimensional equation (31):

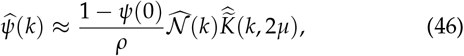

where again we make the approximation that 1 − *ψ*(*x*) is approximately constant over the *x* values where 𝒩(*x*) is non-negligible. This is again accurate for *ψ*(0) ≪ 1, but may need to be adjusted for 1 − *ψ*(0) ≪ 1. While (46) looks exactly like the one-dimensional expression (31), its interpretation is different: *k* is now the magnitude (wavenumber) of the two-dimensional spatial frequency vector (wave vector), *k* = |k|. Note that because dispersal is isotropic, (46) has no angular dependence. The key point is that if we want to transform (46) back to real space, we now must use the two-dimensional inverse Fourier transform. For pairs that are far outside coalescence range, *x* ≫ *δ*, the simple relation (30) between *ψ*(*x*) and 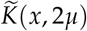 still holds.

#### Lévy flight dispersal

For a two-dimensional Lévy flight, the dispersal kernel of a single lineage takes the form of an isotropic stable distribution (Zolotarev 1981):

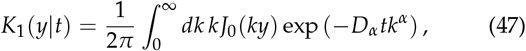

where *K*(*y*| *t*) is the probability density of being at a particular point a distance *y* away from the initial position at time *t*, and *J*_0_ is the zeroth Bessel function of the first kind. As in one dimension, the density for the displacement between the two Lévy flights is the same, but with twice the dispersal constant:

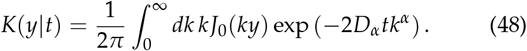

(48) is the two-dimensional inverse Fourier transform (equivalently, the inverse zeroth-order Hankel transform) of the characteristic function 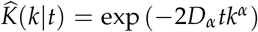. The Fourier-Laplace transform is again 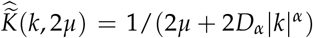. At large distances, *y* ≫ (*D*_*α*_*t*)^1/*α*^, *K* has a power-law tail (Nolan 2013):

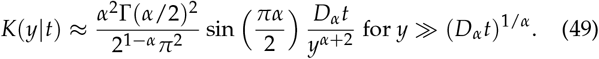

In two dimensions, we must allow coalescence to take place at a finite distance for all *α* (Mörters and Peres 2010). For the coalescence kernel, we use an isotropic normal distribution 𝒩 (*x*) with mean zero and standard deviation *δ*, with coalescence taking place at rate 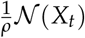. Inverting the Fourier transform in (46) then gives:

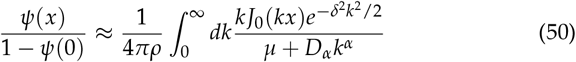

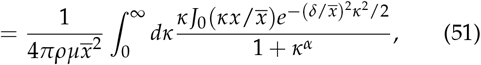

The analysis of (51) parallels that of the one-dimensional case, but all *α* < 2 can be treated together for all distances *x*, not just 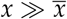, and so we can conduct one unified analysis moving from short distances to long ones.

### Probability of identity for co-located pairs, x = 0

For pairs sampled from the same location, *x* = 0, the Bessel function in (51) is simply equal to one and can be dropped:

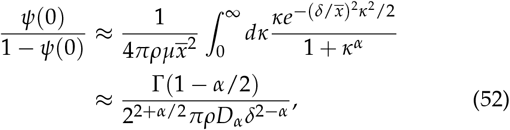

where in the last line we have assumed that 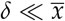. Intuitively, (52) can be understood as roughly the ratio between the time to coalesce, i.e., the neighborhood size ∼ *ρδ*^2^ and the time ∼ *δ*^*α*^ /*D*_*α*_ that the lineages will spend in the same neighborhood before jumping apart. Note that mutation does not enter: in two dimensions, all *α* < 2 act like *α* < 1 does in one dimension, where locally mutation is irrelevant. Again, (52) is only accurate for *ψ*(0) ≪ 1.

Solving (52) for *ψ* gives:

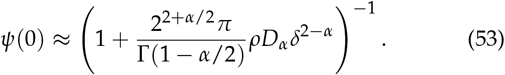

For *α* = 2, integrating (51) with *x* = 0 recovers the classic diffusive result in two dimensions, which we expect to hold for pairs in contact when *α* ≥ 2 (Barton *et al*. 2002):

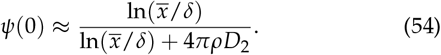

### Probability of identity for separated but nearby pairs, 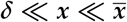

For pairs that are outside coalescence range, *x* ≫ *δ*, we can find *ψ* from (30):

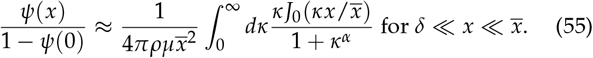

This looks different from the one-dimensional equation (33) because now we had to apply the two-dimensional inverse Fourier transform to 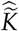 to obtain 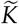. For nearby pairs 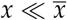, the integral in (55) is dominated by *κ* ≫ 1 and for *α* < 2 we can approximate the denominator in the integrand as 1 + *κ*^*α*^ ≈ *κ*^*α*^, giving:

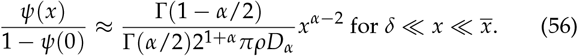

The convergence of (55) to (56) is however quite slow in 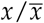 when *α* is close to 0 or 2. For example, for 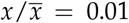, the two expressions differ by ≈ 30 − 40% for *α* = 0.25 and *α* = 1.75, and only approach to within 10% of each other at extreme values of 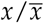 (≈ 10^−5^ and ≈ 10^−4^ for *α* = 0.25 and *α* = 1.75, respectively).

Plugging (53) for *ψ*(0) into (56) lets us solve for *ψ*:

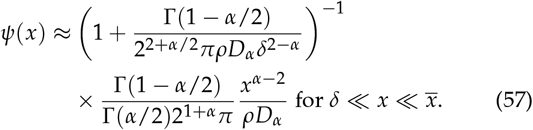

We see that in two dimensions, relatedness at short distances is independent of *μ* to leading order for all *α* < 2. However, the slow convergence mentioned above means that for most biologically reasonable parameter values, this should be interpreted as meaning that the dependence on mutation rate is weak rather than negligible.

For *α* = 1 and 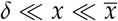 we recover Eq. (A6) of Chave and Leigh Jr (2002) for Cauchy dispersal. Note that they consider distances large compared to the typical single-generation dispersal distance, *c* ≡ (2*D*_*α*_)^1/*α*^, but small compared to 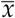, and thus our result for 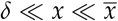 is consistent with their findings.

For *α* = 2 and 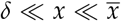 we can recover the known result for diffusive motion by approximating (51) as

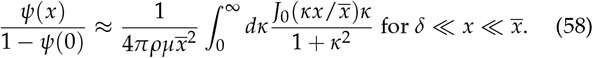

Integrating (58) confirms that we find logarithmic scaling of *ψ*(*x*) at short distances (Barton *et al*. 2002):

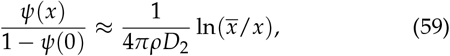

which we expect to hold at 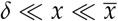 for all *α* ≥ 2.

### Probability of identity by descent for distant pairs, 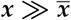

The probability of identity by descent for distant pairs 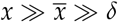 can be immediately be read off from (30) by substituting in the tail of the two-dimensional dispersal kernel (49) for *α* < 2:

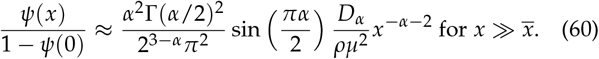

Plugging in (53) for *ψ*(0) lets us solve for *ψ*:

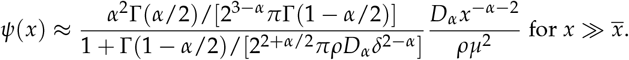

When *α* = 2, we instead recover classic expression for two-dimensional diffusive motion at large distances (Barton *et al*. 2002):

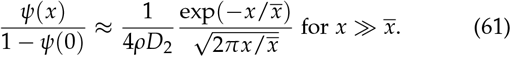

For our simulations, rather than using a true two-dimensional stable distribution, we use radial draws from a one-dimensional stable distribution and then pick a direction at random. The resulting dispersal kernel is shown in (12). At large distances, 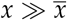, we can apply (37) to find that the tail expression for IBD (when *α* < 2) is simply (*πx*)^−1^ times (36):

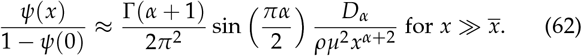

For the finite variance 2D kernel (13) used when *α* > 2, we can again apply (37) to find the tail expression for IBD:

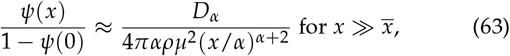

where *D*_*α*_ is defined as *ω*_*α*_ for the single lineage kernel (13).

While the above expressions are accurate in continuous time, we multiply by an extra factor to adjust for discrete time in the tail expressions of Fig. 4 where *μ* = 1. This factor *f* is the ratio between the discrete time sum of *te*^−2*t*^ from *t* = 1 to *t* = ∞ and the continuous time integral of *te*^−2*t*^ from *t* = 0 to *t* = ∞: *f* ≈ 0.7.

### Coalescence time distribution

In this section we will find asymptotic expressions for the coalescence time distribution. As stated in the Results, intuitively we can think of the probability of identity *ψ* as measuring the probability of the pair of lineages coalescing ≲ 1/(2*μ*) generations ago. We can make this statement more rigorous using the Hardy-Littlewood Tauberian theorem connecting the long (short) time probability of coalescence to the small (large) mutation rate limit of *ψ*. It states that a function *f* (*t*) has the limiting behavior 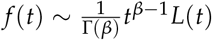 as *t* → ∞ (*t* → 0), where *L* is a slowly varying function and *β* > 0, if and only if its Laplace transform 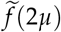 has the limiting behavior 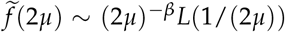 as *μ* → 0 (*μ*→ ∞) (Feller (1971), XIII.5, Theorem 4). Here we follow Feller in using “∼” to denote that the expressions approach each other asymptotically, i.e., that their ratio approaches 1.

### Recent times

First we will consider the limit of recent times, *t* → 0 / *μ* → ∞. For pairs sampled within coalescence range, *x* ≲ *δ*, by definition the coalescence time density approaches *p*(*t* → *x*) ∝ 1/(*ρδ*^*d*^), up to numerical factors that depend on the details of the coalescence kernel. Here *d* is the dimensionality of the range, *d* = 1 or 2. For pairs sampled well outside coalescence range, *x* ≫ *δ*, we can assume that 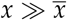 as well, because 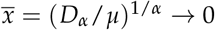 as *μ* → ∞. We can also assume that 1 → *ψ*(0) → 1 is independent of *μ* to leading order. (For *α* < *d* our expressions for *ψ*(0) (43) and (53) are also independent of *μ* and non-zero, but these are only valid when 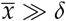, i.e., when *μ* is not arbitrarily large.) We can therefore apply the Tauberian theorem to (36) and (60) to obtain:

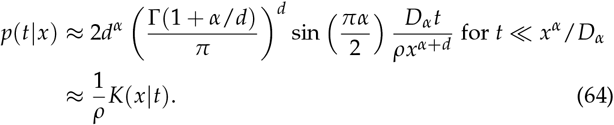

Our heuristic derivation in the Results section essentially proceeded in the opposite direction, starting from 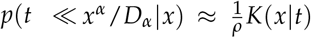 and then deriving *ψ*(*x*) from that. (64) is thus essentially just a restatement of our expressions for the tail of *ψ*, and its accuracy can be seen from the same simulation results shown in Fig. 5.

### Long times

While there is a single unified expression for *p* in the *t* → 0 limit, corresponding to the single expression for *ψ* in the *x* → ∞ limit, for the opposite limit, *t* → ∞ / *μ* → 0, we must treat different values of *α* separately, just as we did for *ψ* at small *x*. We verify our results with simulations, shown in Fig. 7. Note that these expressions will only hold at long times on an infinite range. For any range of finite size, the right tail of the coalescence time distribution will decay exponentially, as in the case of a panmictic population (Wilkins 2004).

For *α* < *d*, we can simply take the inverse Laplace and Fourier transforms of (31) to find *p*, because 1 − *ψ*(0) is independent of *μ* to leading order. Because we are concerned with times long compared to the time for the lineages to traverse the coalescence zone, *t* ≫ *δ*_*α*_ /*D*_*α*_, the normal factor in (31) can be neglected and *p*(*t*|*x*) is simply given by the inverse Laplace transform of (30):

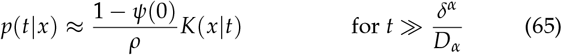

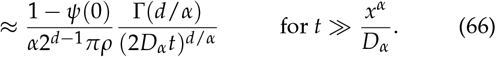

Integrating (66) yields the cumulative distribution for *t* ≫ *x*^*α*^ /*D*_*α*_:

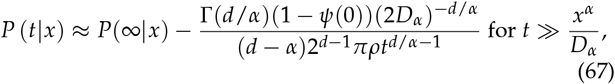

where *P*(∞ | *x*) = lim_*μ* → 0_ *ψ*(*x*) is given by (41), (42), (53), or (57), depending on *x* and *d*.

For *d* ≤ *α* ≤ 2, the leading behavior of the cumulative distribution is trivial: it approaches one at large times, lim_*μ*→ 0_ *ψ*(*x*) = lim_*t* ∞_ *P*(*t* | *x*) = 1. So to find interesting asymptotic behavior, we must instead consider the complementary cumulative distribution, 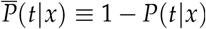, also known as the survival function.

Its Laplace transform is:

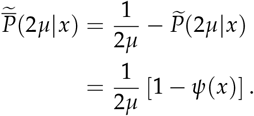

We can now apply the Tauberian theorem to 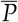 and 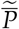. Because we are taking the *μ* → 0 limit, we have 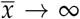, and we need only consider 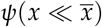.

For *d* = 1 and 1 < *α* ≤ 2, inspecting (39) and (40), we see that they have the limit:

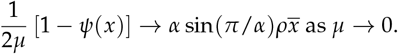

Because 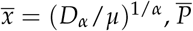 has the limit:

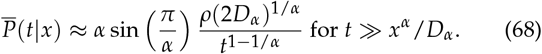

Differentiating (68) yields the density *p*(*t*|*x*):

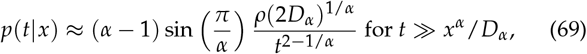

in agreement with Janakiraman (2017)’s Eq. 19.

For *α* = 2, (68) and (69) simplify to the classic diffusive results:

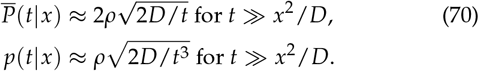

For *α* > 2, we expect the coalescence rate 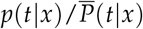 to be-have similarly at long times, as the dispersal approaches a diffusion. But the distribution *P* may be different, due to differences in the probability of early coalescence (Fig. 7, bottom right).

For the marginal cases *α* = *d* = 1 and *α* = *d* = 2, we can find the limit of 1 − *ψ*(*x*) from the expressions for *ψ* in (43), (45), (54), and (59):

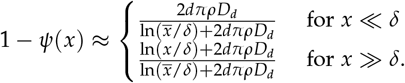

These two limits are similar enough that it makes sense to combine them into one approximation that works for both large and small *x*:

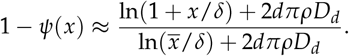

After dividing by 2*μ* and substituting in 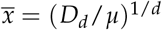, we can then apply the Tauberian theorem to find the complementary cumulative distribution of coalescence times:

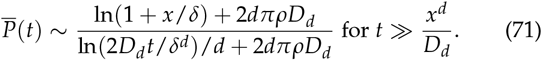

Note that the Tauberian theorem does not guarantee the correctness of sub-leading behavior for *t* → ∞; in this case, that means the term 2*dπρD*_*d*_ in the denominator. However, we have confirmed that is indeed accurate via simulation (Figure 7, *α* = 1, top right). Interestingly, the simulations show that (71) is accurate even for *P*(*t*) ≪ 1. We can also differentiate (71) to find the density *p*:

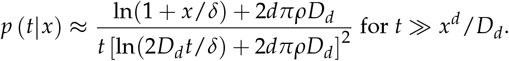

Note that in two dimensions, *α* = *d* = 2 represents the diffusive limit, and we expect these expressions for the marginal case to hold for all *α* ≥ 2.

### Breakdown of models at small scales

Great care must be taken in defining coalescent models in continuous space in order to guarantee that they have a consistent forward-time biological interpretation (Felsenstein 1975; Barton *et al*. 2010). We have not done this, and therefore the microscopic behavior of our continuous approximations does not correspond to any biological population. However, the behavior at large scales (time long compared to one generation, distance long compared to the coalescence scale *δ* and the typical single-generation dispersal distance *c* ≡ (2*D*_*α*_)^1/*α*^) should be accurate (Barton *et al*. 2002). We have shown via simulation that our results accurately describe a stepping-stone model of discrete demes of size ∼ *ρδ*^*d*^ separated by distance ∼ *δ*.

The key place in which the microscopic details matter even for large distances and long times is the factor 1 − *ψ*(0) which appears in many of our expressions. As discussed above, for *α* < *d* even here the microscopic details are not necessarily important, but for *α* ≥ *d* they are. Practically speaking, this quantity would typically have to simply be measured in a population or else treated as a fitting parameter when matching the large-scale predictions to data.

At a microscopic level, we expect that our continuous-time analytic model should deviate from discrete-time models such as the one we use in our simulations. As shown in Fig. 8, this becomes apparent for *α* < 1 in one dimension (or more generally, *α* < *d*). The two differ at scales smaller than the typical single-generation dispersal distance, *x* < *c* = (2*D*_*α*_)^1/*α*^, when this scale is large compared to the coalescence scale, *c* ≫ *δ*. In continuous time, nearby pairs with *x* ≪ *c* would be able to coalesce at times smaller than a single generation, *t* ≪ 1. But in discrete time no pairs can coalesce until *t* = 1, by which time the dispersal kernel *K*(*x*|1) is roughly flat out to *x* ≲ *c*, and probability of identity thus becomes approximately constant for *x*≲ *c*. (For *α* ≥ *d*, the continuous-time model already predicts that *ψ* should be changing slowly at 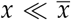, and therefore we do not expect a disagreement with the discrete-time model.) Recall that our discrete-time model assumes no coalescence at *t* = 0 even for lineages starting at *x* < *δ*; if we were to change this, *ψ* would discontinuously jump up to a second, higher plateau for *x* < *δ*.

We can estimate the discrete-time value of *ψ*(*x* ≪ *c*) from a heuristic argument, at least when *ψ* ≪ 1. In the absence of coalescence, the probability of the lineages being within coalescence range of each other in generation *t* ≥ 1 is ≈ (2*δ*)*K*(*x*|*t*) ≈ (2*δ*)*K*(0 | *t*). For *ψ* ≪ 1, including the possibility of coalescence will only slightly decrease this probability. Given that the lineages are in coalescence range, they coalesce with probability 1/(2*δρ*). So in any one generation the probability of coalescence is ≈ *K*(0|*t*)/*ρ* and we can find *ψ* by summing over all generations:

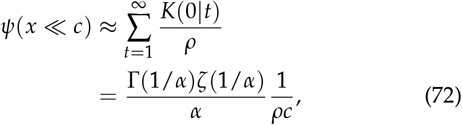

where *ζ* is the Riemann zeta function. Fig. 8 shows that (72) accurately describes the simulations.

**Figure 8.**
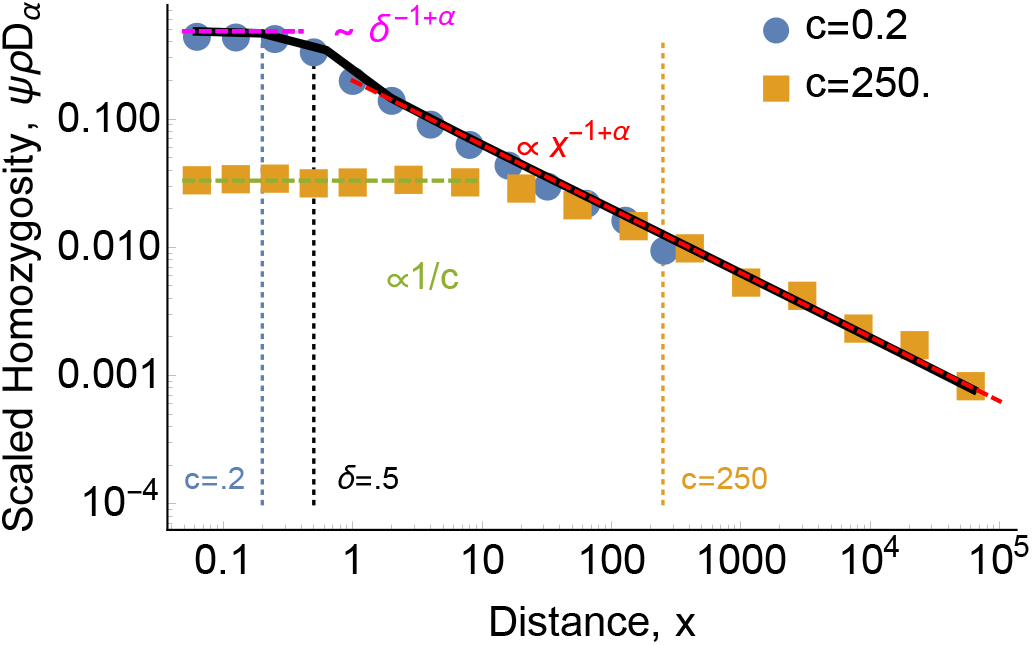
For heavy-tailed dispersal with *α < d*, continuous-time and discrete-time models differ at short distances. Scaled probability of identity *ψ* as a function of distance *x* for *α* = 0.5, *δ* = 0.5, and *ρ* = 100. Points show discrete-time one-dimensional continuous-space simulation results. For the continuous-time model, the black curve shows the result of numerically integrating (33), while the dashed red and magenta lines show the asymptotic approximations (42) and (41), respectively. The continuous-time model predicts that *ψ* should only plateau within the coalescence distance *δ*, but for distance between *δ* and the typical single-generation dispersal distance *c*, the change in *ψ* is driven by the probability of coalescing at 0 < *t* ≪ 1. In the discrete-time model, these lineages have to wait until *t* = 1 to coalesce, leading to a lower, broader plateau, given by (72) (dashed green line). This discrepancy only exists for *δ* < *x* ≪ *c*, i.e., if *c* < *δ* then the discrete-time and continuous-time models agree (blue points).

## Acknowledgments

The authors thank Nick Barton, Shai Carmi, Reed Cartwright, Graham Coop, David Field, and Peter Ralph for helpful suggestions. DBW was supported by a Simons Foundation Investigator award in the Mathematical Modeling of Living Systems, by a Sloan Foundation Research Fellowship, and by NSF CAREER award PHY-2146260. TBS was partially supported by the NSF award 1806833, the iPoLS Student Research Network.

## Appendix A. Alternative derivations of the probability of identity by descent

### Starting from a recursion equation

(31) can also be derived starting from a recursion equation for *ψ* requiring that it remain constant over an infinitesimal timestep *dt* (Malécot 1975; Barton *et al*. 2002):

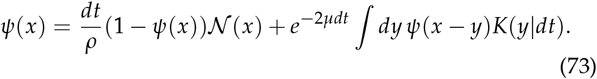

(73) is saying that at equilibrium the local increase in identity due to coalescence (first term) must be balanced by the loss of identity due to mutation and the spreading of identity due to dispersal (both included in the second term).

Taking the spatial Fourier transform ℱ{·} of (73) simplifies the second term at the expense of complicating the first:

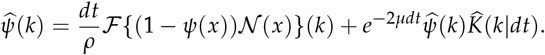

Solving for 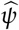 gives:

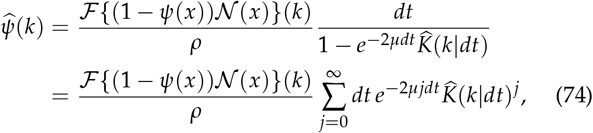

where in the second line we can take the Taylor series expansion because *e*^−2*µdt*^ < 1 and 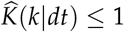 because it is a characteristic function. Assuming dispersal is Markovian, we can simplify (74) by noting that 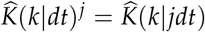, i.e., the distribution after time *jdt* is just the convolution of *j* dispersal steps of time *dt* each. Using this, we can convert the sum into an integral to find (31):

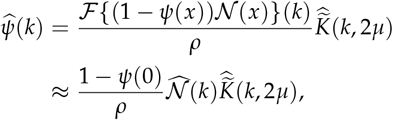

where in the second line we have used the same approximation that 1 − *ψ*(*x*) ≈ 1 − *ψ*(0) for |*x* | ≲ *δ* that we used in the main text.

### Fractional diffusion equation

For Lévy flight dispersal, (32) can also be derived using a fractional diffusion equation. When *X*_*t*_ follows a diffusion, (23) can be written as a Feynman-Kac (diffusion) equation for *ψ* (Barton *et al*. 2002; Allman and Weissman 2018). For *α* < 2, this generalizes to a fractional differential equation:

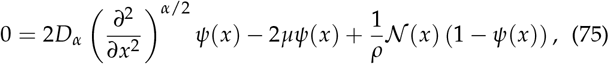

where 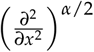 is a Riesz fractional derivative, defined by its Fourier transform 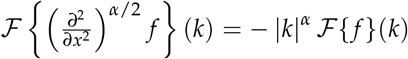 (Metzler *et al*. 2009; Carmi *et al*. 2010; Janakiraman 2017). It is therefore simpler to consider the Fourier transform of (75), which is equivalent to (32):

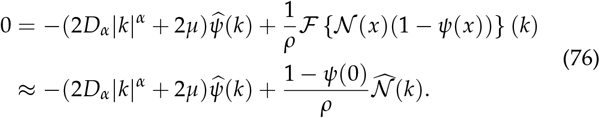

For all *α* < 2, the solution for *ψ* in two dimensions can be also written as a fractional differential equation (Chen *et al*. 2012):

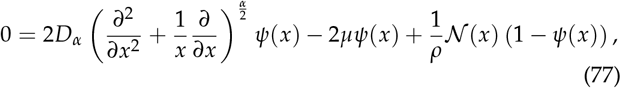

Where 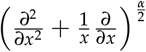 is a fractional Laplacian, defined by its Fourier transform 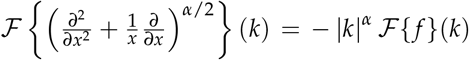 (Kwaśnicki 2017; Lischke *et al*. 2020). Note that the rotational symmetry of the problem allows us to write the Laplacian in terms of just the radial coordinate *x*, and ignore the angular coordinate. The two-dimensional Fourier transform of (77) has exactly the same form as (76), although again the interpretation is different. *k* is now the radial coordinate in two-dimensional *k*-space, i.e., the magnitude of the two-dimensional wavenumber vector.

## Appendix B. Cartoon lineage trajectories in one dimension

In one spatial dimension, the qualitative classification of possible lineage dynamics is similar to that in two dimensions (shown in Fig. 3). But really, the key separation among different forms of power-law dispersal is whether they are recurrent, i.e., the comparison between *α* and the spatial dimension *d*. So while in two dimensions the split is between dispersal with finite or infinite variance, in one dimension it is between dispersal with finite or infinite mean. This is illustrated in Fig. 9.

**Figure 9.**
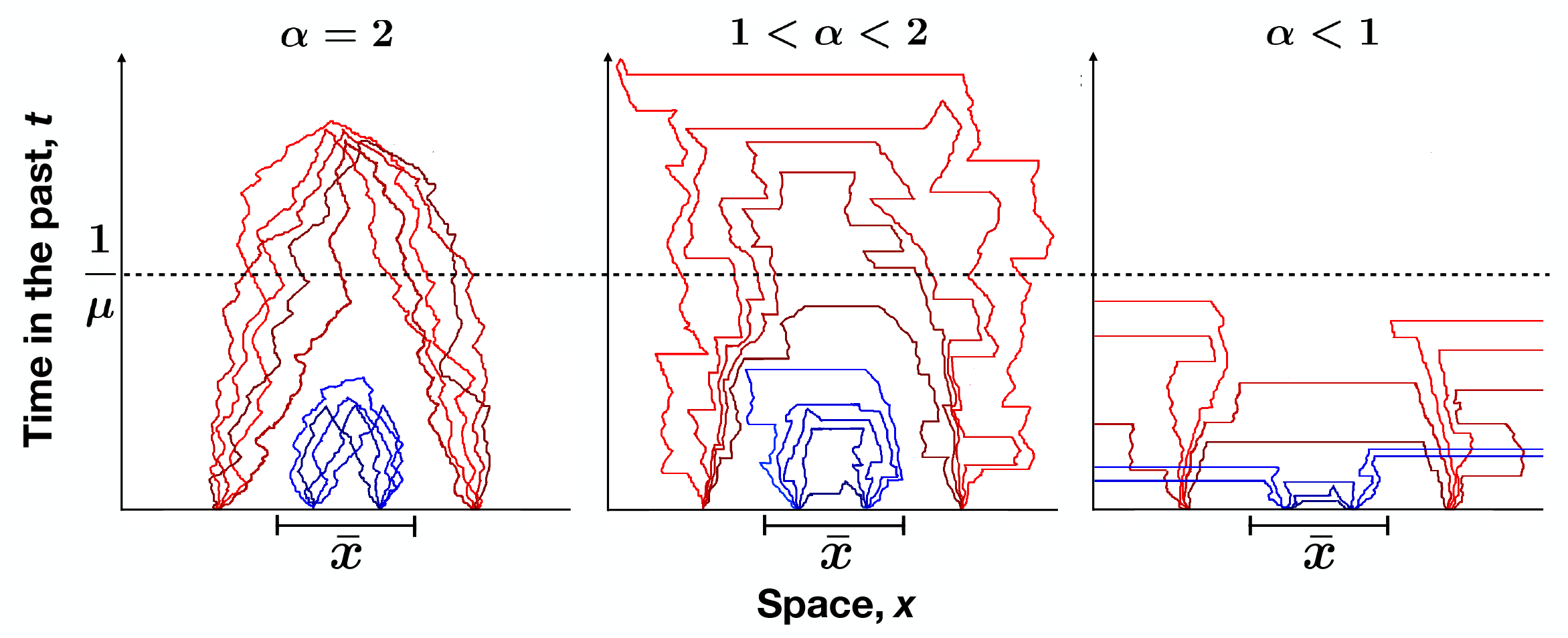
Long-range jumps affect when and where lineages coalesce in one dimension. Qualitative illustrations of lineage dynamics for each of the three *α* regimes of one-dimensional Lévy flights. Typical histories are shown for nearby samples (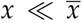, blue) and distant samples (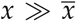, red). Left: For diffusive motion, *α* = 2, the initial separation *x* is a relatively good predictor of coalescence time. Center: For dispersal with an infinite variance but finite mean, 1 < *α* < 2, large jumps broaden the spatial and temporal ranges over which lineages coalesce. Lineages at large separations 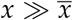 are occasionally able to coalesce at times shorter than 1/*μ*. The coalescence time distribution for nearby lineages 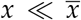 is only modestly affected. Non-Lévy flights with *α* > 1, even those with finite variance (*α* ≥ 2), follow a similar pattern. Right: For dispersal with an infinite mean, *α* < 1, large jumps are common. This allows for rapid coalescence of lineages at both small and large distances, but also lets lineages jump very far away from each other and avoid coalescing until a much later time set by the total range size.

## Notes

https://github.com/weissmanlab/Long_Range_Dispersal

### Competing Interest Statement

The authors have declared no competing interest.

### Summary of Updates

Added exact simulations of a population occupying a finite, two-dimensional lattice of demes.

